# Cell Atlas of Aqueous Humor Outflow Pathways in Eyes of Humans and Four Model Species Provides Insights into Glaucoma Pathogenesis

**DOI:** 10.1101/2020.02.04.933911

**Authors:** Tavé van Zyl, Wenjun Yan, Alexi McAdams, Yi-Rong Peng, Karthik Shekhar, Aviv Regev, Dejan Juric, Joshua R. Sanes

**Author notes:** Equal contribution (Tavé van Zyl, Wenjun Yan). Correspondence to: Joshua R. Sanes or Tavé van Zyl. Y. R. P., Department of Ophthalmology and Jules Stein Eye Institute, UCLA, Los Angeles CA, K.S., Department of Chemical Engineering, UC Berkeley, Berkeley, CA.

## Abstract

Increased intraocular pressure (IOP) represents a major risk factor for glaucoma, a prevalent eye disease characterized by death of retinal ganglion cells that carry information from the eye to the brain; lowering IOP is the only proven treatment strategy to delay disease progression. The main determinant of IOP is the equilibrium between production and drainage of aqueous humor, with compromised drainage generally viewed as the primary contributor to dangerous IOP elevations. Drainage occurs through two pathways in the anterior segment of the eye, called conventional and uveoscleral. To gain insights into the cell types that comprise these pathways, we used high-throughput single cell RNA sequencing (scRNA-seq). From ∼24,000 single cell transcriptomes, we identified 19 cell types with molecular markers for each and used histological methods to localize each type. We then performed similar analyses on four organisms used for experimental studies of IOP dynamics and glaucoma: cynomolgus macaque (*Macaca fascicularis*), rhesus macaque (*Macaca mulatta*), pig (*Sus scrofa*) and mouse (*Mus musculus*). Many human cell types had counterparts in these models, but differences in cell types and gene expression were evident. Finally, we identified the cell types that express genes implicated in glaucoma in all five species. Together, our results provide foundations for investigating the pathogenesis of glaucoma, and for using model systems to assess mechanisms and potential interventions.

## INTRODUCTION

Glaucoma, the leading cause of irreversible blindness worldwide (Quigley and Bromann, 2006), results from loss of retinal ganglion cells (RGCs), which carry information about the visual world from the eye to the rest of the brain (Weinreb et al., 2014). Of the major risk factors, including age, race and family history, the only modifiable one is intraocular pressure (IOP). Indeed, lowering IOP remains the only proven treatment strategy to delay disease progression, with several pharmacological and surgical approaches in widespread clinical use. However, therapeutic advances have been limited by incomplete understanding of the tissues that regulate IOP and the means by which increased IOP leads to RGC loss. We address the first of these issues here.

There are two major compartments in the eye, an anterior segment containing aqueous humor (AH) and a posterior segment containing vitreous humor. The primary determinant of IOP is the equilibrium between the production and drainage of AH. AH is produced by the ciliary body, circulates within the anterior chamber, and is then drained through one of two pathways, conventional (trabecular) or uveoscleral (Costagliola et al., 2019) (***Figure 1A,B***). In the conventional pathway, AH exits the anterior chamber through a lattice-like biological filter called the trabecular meshwork (TM), which is composed of collagenous beams lined with specialized TM cells. It then passes through a region of cells embedded within a denser extracellular matrix called the juxtacanalicular tissue (JCT) prior to being conveyed into a specialized vessel called Schlemm canal (SC). From SC, AH exits the eye through a network of collector channels (CC) continuous with the venous system. The remaining AH exits the anterior chamber via the uveoscleral pathway, draining through the interstices of the ciliary muscle, ultimately exiting the eye via the suprachoroidal space and the sclera. In principle, increased IOP could result from either excessive production or insufficient drainage of AH, but in practice, the latter is more commonly implicated (Grant 1958; Braunger et al., 2015; Stamer and Clark, 2016).

**Figure 1.**
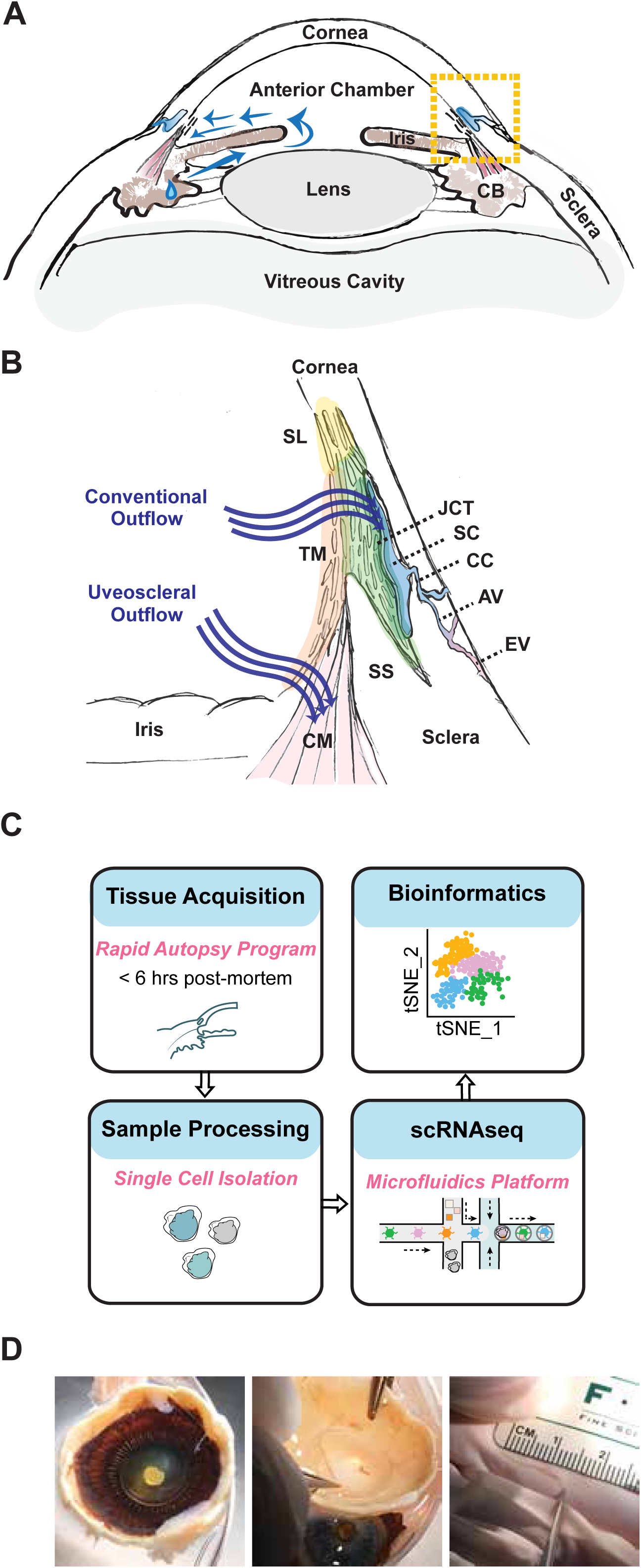
Human aqueous humor outflow pathways. **A** Diagram of the anterior segment of the human eye, which includes the cornea, iris, ciliary body and lens. Aqueous humor is secreted by the ciliary body (CB) and circulates (blue arrows) within the anterior chamber prior to draining from the eye through one of two pathways located within the iridocorneal angle, delineated here by the boxed area. **B** Enlarged view of the iridocorneal angle, boxed area in (A), highlighting two outflow pathways for aqueous humor (AH). In the conventional pathway, AH traverses the trabecular meshwork (TM), first through the uveal meshwork (orange highlight), then the corneoscleral meshwork (light green highlight) and finally the juxtacanalicular tissue (JCT, dark green highlight) prior to entering Schlemm canal (SC). AH exits the SC via collector channels (CC) that empty into aqueous veins (AV) that themselves merge with episcleral veins (EV). Non-filtering TM is located at the insert region (yellow highlight), referred to as Schwalbe line (SL), which abuts the corneal endothelium. In the uveoscleral pathway, AH exits via the interstices of the ciliary muscle (CM). SS, Scleral spur. **C** Workflow for obtaining single cell transcriptomes. **D** Dissection procedure. Left: anterior segment following dissection at the pars plana, posterior view. Middle: blunt dissection of TM strip following removal of lens, iris and ciliary body. Right: Strip of isolated TM.

The cellular composition of the AH outflow pathways has been studied histologically (Alvarado et al., 1984; Tamm, 2009), but because of their small size, intricate architecture and tightly invested tissues, it is impractical to dissect individual components for molecular analysis. Therefore, most studies of gene expression in these tissues have relied on bulk measurements of tissue or cultured cells (Liu et al., 2013; Sathiyanathan et al., 2017; Carnes et al., 2018). It has therefore been difficult to assess gene expression patterns of individual cell types. To circumvent this limitation, we used high throughput single cell RNA sequencing (scRNAseq; ***Figure 1C***). We profiled ∼24,000 single cells from adult human TM and surrounding tissues, applied computational methods to cluster the cells based on their transcriptomes, and used histological methods to match the molecularly defined clusters to specific cell types. We identified 19 cell types and defined molecular markers for each of them.

We then used this cell atlas in two ways. First, we assessed expression in each cell type of genes that have been implicated in glaucoma, either as causal genes with Mendelian inheritance or as susceptibility loci identified in genome-wide association studies (GWAS) (Wiggs and Pasquale, 2017; Lewis et al.,2017; Choquet et al., 2018; Gao et al., 2018; Khawaja et al., 2018; Macgregor et al., 2018; Sears et al., 2019; Youngblood et al., 2019; Krumbiegel et al.,2019), and compared expression levels in cell types of the outflow pathways to those in retinal RGCs and retinal glia (Yan et al., in prep). We found that genes associated with elevated IOP were more likely to be preferentially expressed in the anterior segment, whereas those associated with normal tension glaucoma were more likely to be expressed predominantly in the retina.

Second, we used the human atlas as a foundation for assessing four animal models frequently used to study AH outflow pathways and glaucoma – Cynomolgus macaque (*Macaca fascicularis*), rhesus macaque (*Macaca mulatta*), pig (*Sus scrofa*) and mouse (*Mus musculus*) (Bachmann et al., 2006; Fernandes et al., 2015; Picaud et al.,2019). The utility of these models depends in large part on the extent to which their cell types and patterns of gene expression in them correspond to those in humans. We show that broad cell classes were generally conserved across all five species and could be localized to expected areas within the eye, but that some cell types within classes were less well conserved. In some cases, molecular markers specific for human cell types were either absent or much less specific for cell types within other species. Disease genes in humans, which often demonstrated highly specific expression in outflow pathway cell types, mapped well in macaque species but less reliably in the pig and mouse. These results may help guide tests of therapeutic strategies in these models.

## RESULTS

### Cell atlas of human trabecular meshwork and aqueous outflow structures

We used a droplet-based method (Zheng et al., 2017) to obtain 24,023 high quality single cell transcriptomes from human trabecular meshwork tissue dissected from 7 eyes of 6 post-mortem donors (***Figure 1D and Table S1***). Computational analysis (see Methods) divided these cells into 19 clusters (***Figure 2A)***, ranging in frequency from 0.1-36% of cells profiled ***(Figure 2C)***. Cells in nearly all clusters were obtained from all individuals (***Figure 2B***; C7 and C9 discussed below). We identified selectively expressed genes for each cluster (***Figure 2D***), then used *in situ* hybridization and immunohistochemistry to relate the molecularly defined clusters to cell types defined by classical criteria of morphology and location.

**Figure 2.**
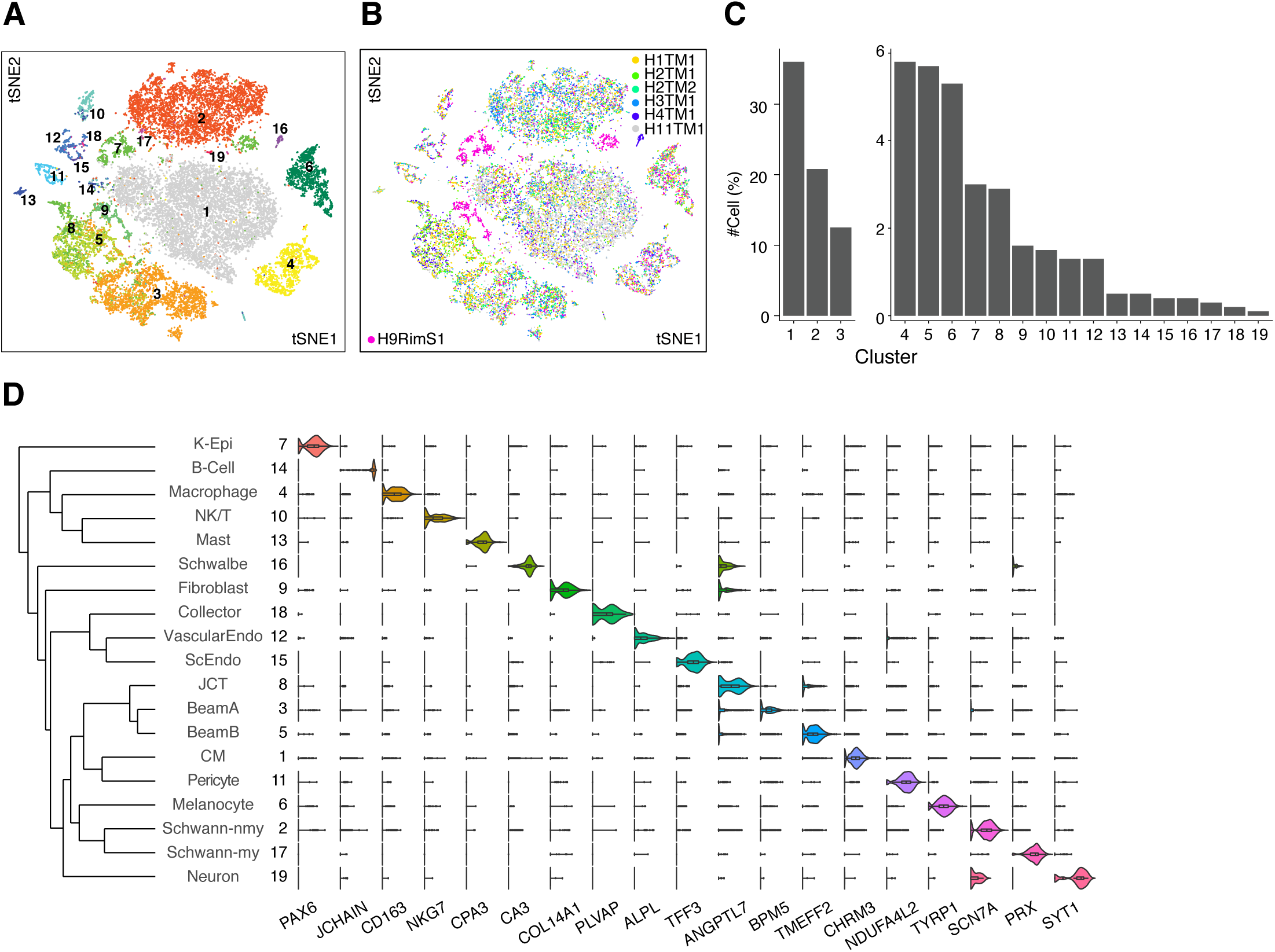
Cell types and gene expression by cells of the human outflow pathways. **A** Clustering of 24,023 single-cell expression profiles from human trabecular meshwork and associated structures visualized by t-distributed stochastic neighbor embedding (t-SNE). Arbitrary colors are used to distinguish clusters deemed to be distinct by unsupervised analysis. Clusters were numbered according to relative size, with 1 being the largest. **B** tSNE plot shown in Figure 2A, but with cells colored by sample of origin. Note that corneal epithelial cells (C7) and fibroblasts (C9) were derived primarily from the rim sample (H9). **C** Frequency of each cell type; numbering as in **A** and **D** **D** Violin plots showing expression of genes selectively expressed by cells of each type. Dendrogram on left shows transcriptional relationships among cell types. K-Epi, corneal epithelium; JCT, juxtacanalicular tissue; CM, ciliary muscle; Mø, macrophage.

#### Conventional pathway

The conventional pathway includes the trabecular meshwork, Schlemm canal, collector channels and closely associated structures **(*Figure 1B*)**. The trabecular meshwork includes an anterior, non-filtering, insert region, and a more posterior filtering region, which itself is divided into two parts: an inner part comprised of “beam cells” through which AH first filters, and an outer part comprised of JCT cells residing adjacent to Schlemm canal, which modulates outflow resistance through the generation and degradation of extracellular matrix (Acott and Kelley, 2008; Stamer and Clark, 2017).

We identified 8 cell types within the conventional pathway, 3 of which corresponded to filtering trabecular meshwork cells. They were characterized and localized as follows:

Clusters 3, 5 and 8 demonstrated high expression levels of *MYOC, MGP*, and PDPN (***Figure 3A***), markers previously associated with trabecular meshwork, as well as other genes such as *RARRES1* (Birke et al., 2010; Watanabe et al., 2010; Stamer and Clark, 2017). We used immunostaining for *PDPN* and *RARRES1* to show that these genes were indeed expressed by cells encasing the trabecular beams (***Figure 3B-D***). Clusters 3 and 5 could be distinguished by preferential expression of *FABP4 and TMEFF2*, respectively, each of which marked a subset of beam cells, which we call Beam A and Beam B (***Figure 3H and S1A, J***). The *FABP4+* Beam A cells and TMEFF2+ Beam B cells were intermingled, but with a tendency for the latter to be closer to the JCT. Cluster 8 could be distinguished from C3 and C5 by selective expression of *CHI3L1* and *ANGPTL7*, both previously suggested to be potential markers for a TM cell subpopulation (Liton et al., 2006). *In situ* hybridization demonstrated localization of cells expressing these genes predominantly to the JCT (***Figure 3I and S2I***). Other DE genes in this cluster (C8) included *RSPO4, FMOD, and NELL2*. Thus, our transcriptomic method identifies three cell types comprising the filtering TM: two types of beam cells and a distinct JCT cell.

**Figure 3.**
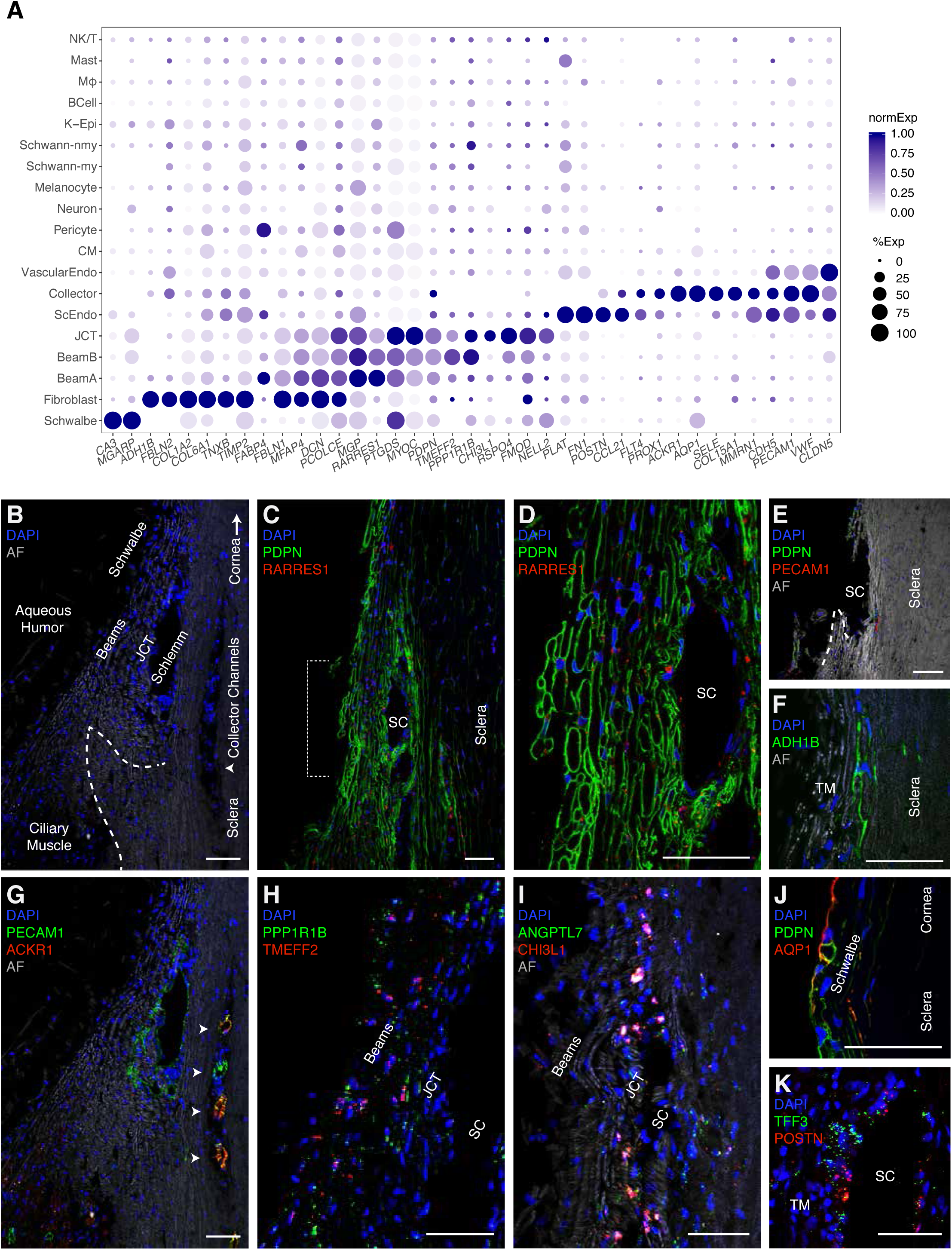
Cells of the Human Conventional Outflow Pathway. **A** Dot plot showing genes selectively expressed in cells of the conventional outflow pathway. In this and subsequent figures, the size of each circle is proportional to the percentage of cells expressing the gene, the color depicts the average normalized transcript count in expressing cells and abbreviations are as in **Fig. 1D**. **B** Labeled meridional section of a human corneoscleral rim visualized using autofluorescence, demonstrates outflow anatomy at the iridocorneal angle visualized after removal of iris and ciliary body. Dashed line indicates scleral spur. **C** Trabecular meshwork (beam & JCT) cells immunostained with *PDPN* (green) and *RARRES1* (red). SC, Schlemm’s canal **D** Higher magnification of region bracketed in **C**. **E** Corneoscleral rim section demonstrates the tissue left behind after TM dissection. No cells in this region are positive for *PDPN* (green) or *PECAM1/CD31* (red), indicating successful removal of relevant structures (TM and SC) during dissection protocol. **F** Scleral fibroblasts identified in the corneoscleral rim collection immunostained with*ADH1B* **G** Immunostaining against *PECAM1/CD31* (green) highlights Schlemm canal, collector channels, and ciliary muscle capillaries, while *ACKR1/DARC* co-stains only the collector channels. Same field as **B**. **H** Fluorescent RNA *in situ* hybridization against *TMEFF2* (green) and *PPPR1B1* (red) highlights beam cells. **I** Fluorescent RNA *in situ* hybridization against *ANGPTL7* (green) and *CHI3L1* (red) highlights cells in the JCT. **J** Schwalbe line cells at the junction of TM and corneal endothelium co-stain with PDPN (green) and *AQP1* (red). **K** Fluorescent RNA *in situ* hybridization against POSTN and TFF3 highlights Schlemm canal endothelium. Bars show 50µm.

A fourth cluster (C16) corresponded to cells of Schwalbe’s line, which is located in the “insert” region of the anterior non-filtering meshwork abutting the peripheral corneal endothelium. This cluster had DE genes associated with corneal endothelium (e.g., *CA3, MGARP, SLC11A2, TGFBI*), but some cells shared markers associated with beam cells (e.g., *MYOC, IGFBP2, NELL2, PTGDS*). The candidate TM marker *AQP1* was preferentially expressed in these non-filtering TM cells, consistent with the observation that deletion of *AQP1* in mice did not affect outflow facility (***Figure 3J***) (Zhang et al., 2002). Although it has been suggested that Schwalbe’s line contains TM precursors (Kelley et al., 2009), we detected no DE genes suggestive of “stemness” (Yun et al., 2016) in this cluster; if present, they may have been too rare to be detected.

Schlemm canal endothelial cells (C13) expressed canonical endothelial markers *PECAM-1* (CD31), VE-cadherin (*CDH5*), and claudin-5 (*CLDN5*) as well as the hemostasis genes *PLAT* (tPA) and *VWF* ***(Figure 3A)***. This cluster also selectively expressed the lymphatic endothelial cell markers *CCL21* (secondary lymphoid-tissue chemokine) and *FLT4* (*VEGFR3*), consistent with the notion that SC is a modified lymphatic vessel (Aspelund et al., 2014; Thomson et al., 2014; Ulvmar and Makainin, 2016). Other lymphatic endothelial markers expressed in SC at lower levels were *PROX-1* and *LYVE-1*. Fibronectin-1 (*FN1*), a TGF-beta inducible ECM gene implicated in glaucoma pathogenesis, was differentially expressed in SC endothelium, as were *MMRN1* and *PLVAP* (Herrnberger et al., 2012; Medina-Ortiz et al., 2013). We used the markers *POSTN* and *TFF3* (***Figure 3K***) to distinguish endothelium of the outflow tract from the vascular endothelium in the ciliary muscle, which shared many endothelial markers (discussed later). Some intermingled cells in Schlemm canal expressed CD31 and the vascular endothelial marker *ALPL* (***Figure 4C***) raising the possibility that this structure contains two cell types.

**Figure 4.**
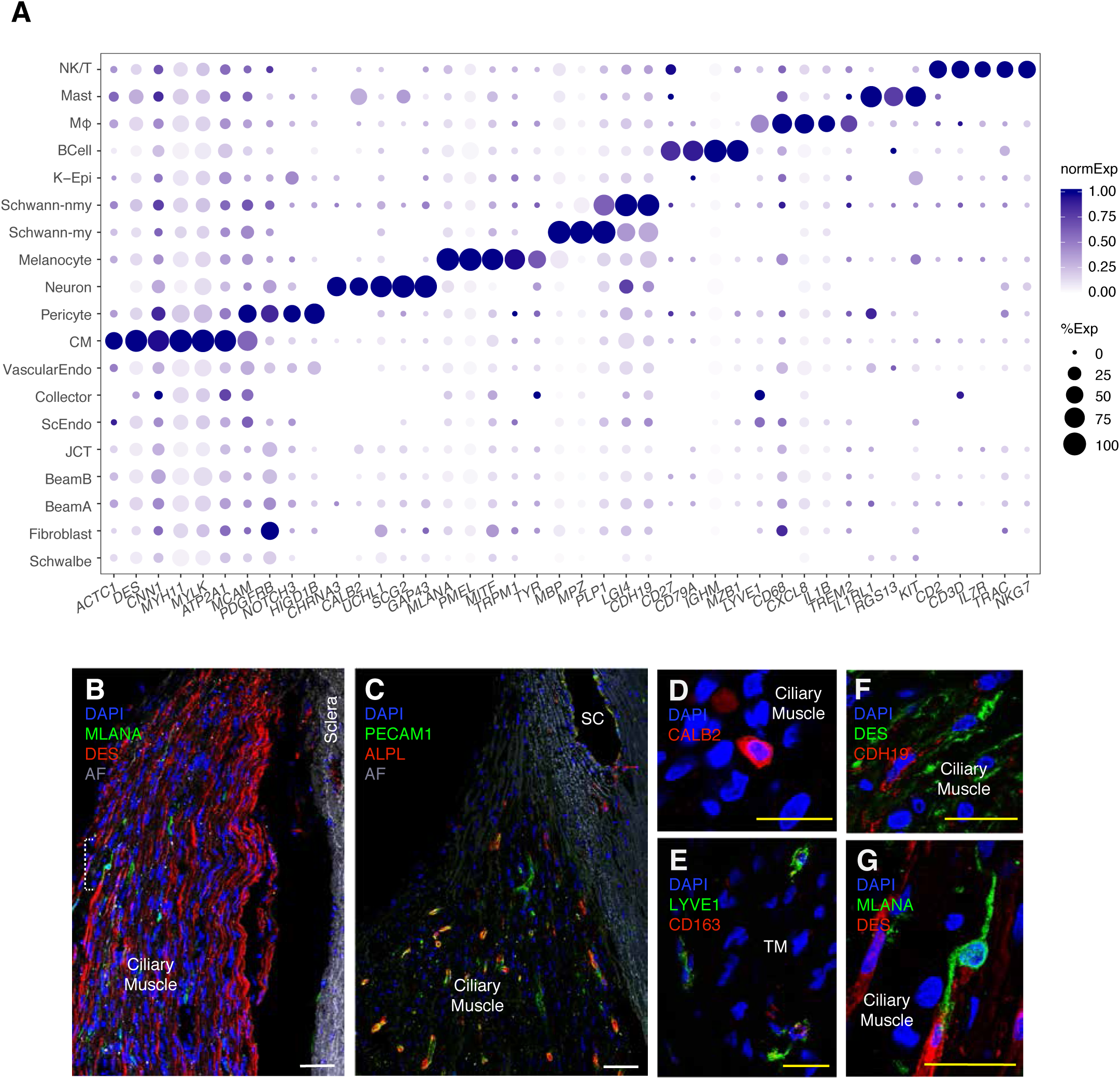
Cells of the Human Uveoscleral Pathway. **A** Dot plot showing genes selectively expressed in cells of the uveoscleral outflow pathway. **B** Smooth muscle cells immunostained with *DES* (red) and melanocytes stained with *MLANA* (green) in ciliary muscle. **C** Capillaries in the ciliary muscle immunostained with *PECAM1* (green) and ALPL (red). Occasional *PECAM1*+*ALPL+* staining was also noted in Schlemm Canal (SC) suggesting that this structure contains more than one cell type. **D** Immunostaining against *LYVE1* (green) and *CD27* (red) identify macrophages in the TM. **E** Immunostaining against *CALB2* (red) highlight intrinsic neurons of the ciliary muscle. **F** Schwann cells in the ciliary muscle stained with *CDH19* (red), amidst ciliary muscle cells stained for desmin (DES; green). **G** Higher magnification of area bracketed in **B** demonstrates MLANA-positive melanocyte (green). Bars show 50µm in B-C, and 25µm in D-G.

Because blunt dissection of the TM may have left behind components of the conventional outflow structures such as the outer wall of Schlemm canal, we collected a corneoscleral rim sample, which included all adjacent tissues after blunt dissection (***Figure 3E***). This sample contained an additional vascular cell type (C19) and a fibroblastic type (C9) that were poorly represented in the other samples. Using the differentially expressed gene *ACKR1* (aka DARC), we identified C19 as the endothelium of the collector channels downstream from Schlemm Canal (***Figure 3G***). *ACKR1* has been identified as a marker to distinguish venular from non-venular endothelial cells, which is consistent with the pre/perivenular location of the channels (Thiriot et al.,2017). Other DE genes for collector channel cells include *SELE, SELP, COL15A1*. The other new cluster, C9, consisted of matrix fibroblasts. Immunostaining for the selectively expressed gene *ADH1B* showed that C9 cells were present within the sclera located adjacent to the TM and outer wall of Schlemm canal (***Figure 3F and S2H***). This cluster was distinguished from TM cell types through its higher level of expression of *FBLN2, TIMP2, TNXB*, and multiple collagen genes (*COL1A2, COL6A1, COL6A2, COL6A3, COL14A1*), markers consistent with previous single cell studies on fibroblasts in other organ systems (Xie et al., 2018). However, it also shared a set of ECM and complement genes with TM cells (C3,5,8) including *DCN, PCOLCE, FBLN1, MFAP4, SERPING1, C1S*, and *C1R*, suggesting that these neighboring cells play overlapping roles.

Finally, the corneoscleral rim sample included cells with transcriptomic signatures marking them as corneal epithelium based on expression of previously described markers such as *AQP3, AQP5* and *KRT5* (Diehn et al., 2005). These cells will be described elsewhere.

#### Uveoscleral pathway

AH that does not exit the eye through the conventional pathway instead exits via the uveoscleral pathway, draining through the interstices of the ciliary muscle. Seven clusters in our dataset comprised components of this pathway. The largest cluster (C1) corresponded to ciliary muscle cells (Wiederholt et al., 2000). It expressed markers classically associated with well-differentiated smooth muscle, including *DES, CNN1, MYH11, MYLK and ACTC1* (***Figure 4A***). We used the DE genes *DES* and *CHRM3* to localize this cluster histologically to the longitudinal ciliary muscle (***Figure 4B and S1B***). The high expression levels of *ATP2A1/SERCA1*, a marker of type II fast twitch skeletal muscle, suggest that ciliary muscle may also have characteristics of skeletal muscle, although it lacks expression of other classical skeletal muscle markers including *MYH1*,-*2*,-*4* and -*7*.

Six additional cell types were found within the ciliary muscle. One, vascular endothelium (C12), was associated with intramuscular capillaries. This cluster shared canonical endothelial markers with Schlemm canal and collector channels but was distinguished from the latter by the presence of *ALPL* (***Figure 4C***) and absence of lymphatic markers. Similarly, PLVAP, while present in Schlemm canal and collector channels, was not expressed in ciliary muscle capillary endothelium, consistent with observations that these cells do not have pores (Ishikawa 1962).

C11 comprises contractile pericytes, which express canonical markers *PDGFRB, MCAM/CD146*, and *NOTCH3* as well as *HIGD1B* (Barron et al., 2016). These cells were labeled by immunostaining against *NDUFA4L2*, and were found within the ciliary muscle and also wrapped around small vessels (***Figure S1C***).

A small neuronal cluster expressing *CHRNA3, UCHL1, SCG2*, and *GAP43* was identified and localized to the ciliary muscle with immunostaining against *CALB2* (aka calretinin) and *ELAVL4 (*aka *HuD)* ***(Figure 4D, S1D)***. These cells presumably correspond to the sparse neurons described in histological and ultrastructural studies (Tamm et al., 1995; Flugel-Koch et al., 2009).

Two clusters, C2 and C17, expressed markers characteristic of Schwann cells, consistent with prior electron microscopy studies showing that Schwann cells ensheath axons within ciliary muscle (Ishikawa 1962). Markers common to both clusters included *PLP1* and *LGI4*. They were distinguished by differential expression of *CDH19* in C2 and a set of genes that encode myelin components, such as *MBP, MPZ*, and *PMP2*, in C17. Immunostaining for CDH19 marked these cells within the ciliary muscle **(*Figure 4F*)**. We conclude that C2 and C17 represent non-myelinating and myelinating Schwann cells, respectively.

Finally, C6 comprised uveal melanocytes, demonstrating expression of canonical markers *MLANA, PMEL, MITF, TRPM1* and *TYR*. Using the marker *MLANA*, we localized these cells to the ciliary muscle ***(Figure 4G)***.

#### Immune cells

Our dataset also included 4 types of immune cells: B cells, NK/T cells, mast cells and macrophages. The macrophages (C4) were *CD163*+ and *LYVE1*+ (Margeta et al., 2016) and localized predominantly to the trabecular meshwork ***(Figure 4E)***. They also expressed *CD68, CD14, CCL3, CCL4, CXCL8, IL1B, TREM2* and *MS4A* genes, all of which have been associated with macrophages in other tissues. Mast cells were localized to the TM using the marker *IL1RL1* and also expressed *CPA3, RGS13*, and *KIT* (***Figure S1E***). B cells, characterized by expression of *CD27, CD79A, IGHM, IGKC, MZB1* and *JCHAIN* were found in only one donor sample but were identified histologically in tissues from other donors using the marker *CD27* (***Figure S1F***). NK/T cells were identified by expression of the DE genes *CD2, CD3D, IL7R, TRAC, GZMA, GZMB*, and *NKG7*.

### Cell-type specific expression patterns of glaucoma-associated disease genes

To improve understanding of how genes associated with glaucoma contribute to disease pathogenesis, we mapped their expression by the 19 cell types in our atlas. We included both known monogenic causes (Mendelian genes) and genes implicated as risk factors in GWAS studies (Wiggs and Pasquale, 2017; Lewis et al.,2017; Choquet et al., 2018; Gao et al., 2018; Khawaja et al., 2018; Macgregor et al., 2018; Sears et al., 2019; Youngblood et al., 2019; Krumbiegel et al.,2019). Mendelian genes assessed were *ANGPT1, ANGPT2, CPAMD8, CYP1B1, FOXC1, LOXL1, LTBP2, MYOC, OPTN, PITX2, TEK* (*TIE2*), and *TBK1*. Well-established GWAS loci genes include *ARHGEF12, ATXN2, CAV1, CAV2, TXNRD2, TMCO1*. We also screened an additional 462 genes listed in the NHGRI-EBI GWAS Catalog for notable expression patterns (Buniello et al., 2019). Overall, 189 genes were expressed in a minimum of 10% of cells in any given class at a minimum expression level of 0.5 log (TPM+1). Examples are shown in ***Figure 5*** with a full list in ***Figure S2***.

**Figure 5.**
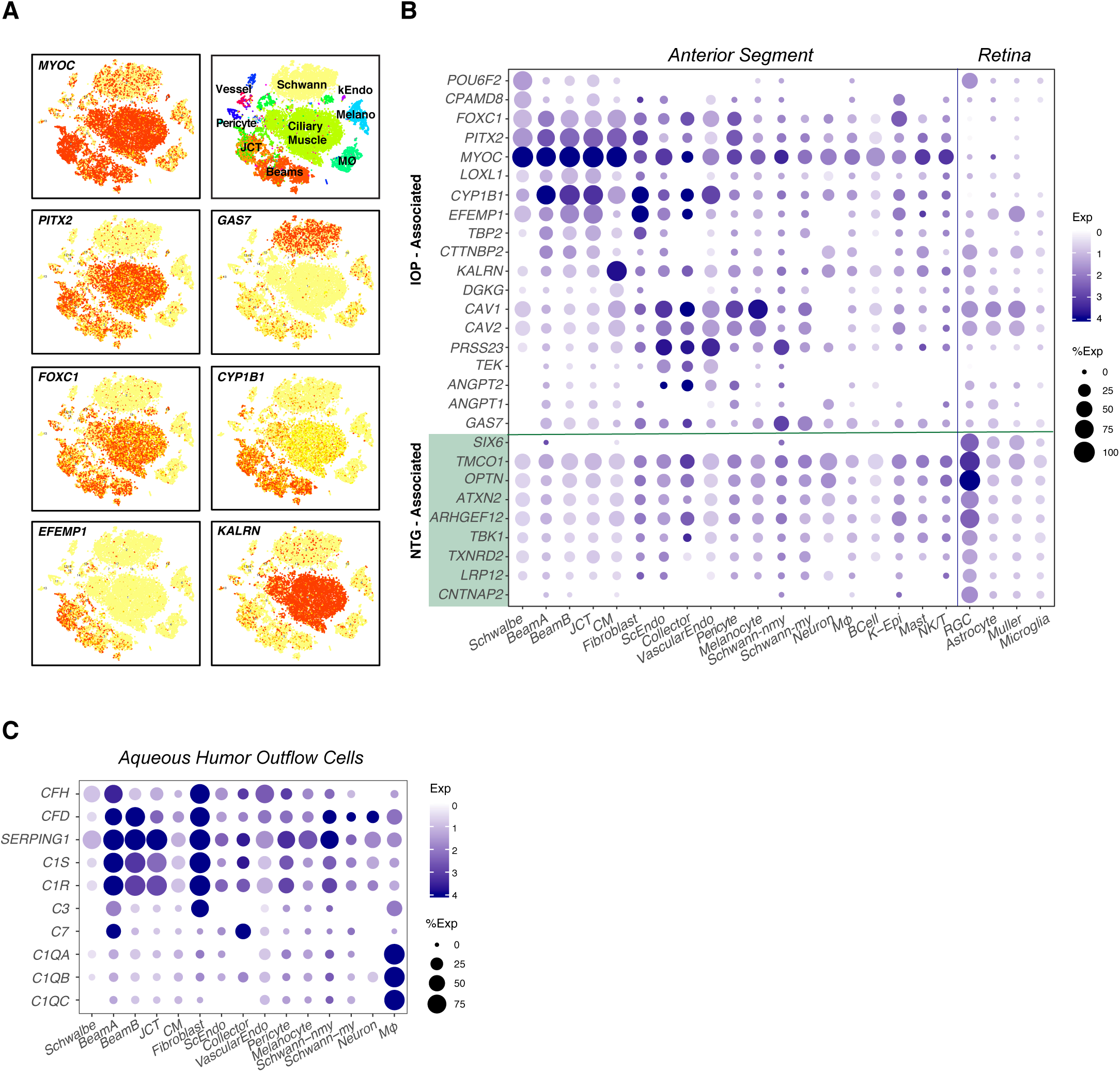
Human disease genes. **A,B**. Cell-type specific expression of several genes implicated in glaucoma, as illustrated by tSNE (A; arranged as in **Fig. 2A**) and dot plot (B). **B** also includes data from a retinal cell atlas, showing expression in RGCs and three types of retinal glia. NTG, Normal Tension Glaucoma. **C**. Dot plot showing cell-type specific expression of complement genes.

Most genes implicated in juvenile glaucoma and high IOP (e.g. *MYOC, FOXC1, PITX2, CYP1B1*), were expressed strongly by all three TM cell types (Beam A, Beam B and JCT), with no evidence for selective expression by any one of the three ***(Figure 5A,B)***. Notably, they were all also expressed at high levels in ciliary muscle. *EFEMP1* exhibited robust cell-type specific expression in beam cells and JCT; in contrast to the first group, it was not present in ciliary muscle, and was expressed differentially among the TM types, JCT>Beam B>Beam A. In contrast, some genes demonstrated stronger expression in non-TM cell types than in TM, including *GAS7* (ciliary muscle, Schwann cells), *KALRN* (ciliary muscle), *PRSS23* (Schlemm canal, collector channel, vessel endothelium and Schwann cells), and *CAV1/CAV2* (all three endothelial types, ciliary muscle, pericytes and melanocytes). Thus, although defects in the TM are clear contributors to elevated IOP, genes that regulate IOP may not act exclusively within the TM.

Because glaucoma risk involves susceptibility of RGCs to degeneration as well as increased IOP, we also interrogated expression in four relevant retinal cell types: RGCs and three glial types with which they interact, astrocytes, Müller glia and microglia **(*Figure 4B*)**. Data for retinal cells are taken from a human retinal cell atlas (see Methods and Yan et al., in prep). We found a distinction between genes associated with primary open angle glaucoma (POAG) involving elevated IOP, and those with Normal Tension Glaucoma (NTG), in which IOP is not elevated. Whereas most genes associated with elevated IOP were expressed in cells of the outflow pathways, as discussed above, genes more closely associated with NTG, including *OPTN, ATXN2, TMCO1* and *SIX6* were expressed at highest levels in RGCs.

Several susceptibility genes, including *CAV1, CAV2* and *POU6F2*, were expressed at high levels in both anterior segment and retina. *POU6F2* is a transcription factor that has been linked to thinner than average central corneal thickness, a highly heritable trait and also a strong risk factor for the development of POAG (Gordon et al., 2002). The expression of *POU6F2* has been detected in human RCGs (Zhou et al., 1996), mouse corneal limbal stem cells, and a subset of mouse RGCs (King et al., 2018). We found preferential expression of *POU6F2* in both corneal endothelium (specifically, Schwalbe line cells) and RGCs ***(Figure 5B)***.

Two genes associated with exfoliation glaucoma (XFG) and the related exfoliation syndrome (XFS), *LOXL1* and *CNTNAP2*, demonstrated divergent regional expression patterns: the former localized predominantly to TM (beam cells and JCT), whereas the latter localized to RGCs. This result offers further insight into XFG as both a primary and secondary open angle glaucoma, characterized not only by IOP elevation due to TM obstruction but also an inherent susceptibility to glaucomatous optic neuropathy.

Finally, motivated by extensive literature implicating the complement system in ocular disease pathogenesis and progression (Clark and Bishop, 2018; Pauly et al., 2019), we investigated the expression of complement genes in the anterior segment. We found many genes encoding complement factors were selectively expressed in the conventional outflow pathway ***(Figure 5C)***. For example, expression of C1qa, C1qb and C1qc, was observed in resident macrophages; the serine proteases C1R and C1S, the C1-inhibitor SERPING1, and other complement genes CFD, CFH, C3 and C7 were expressed preferentially in both TM cells and scleral fibroblasts.

### Model Species

Analysis of mechanisms underlying IOP regulation and tests of therapeutic interventions rely almost entirely on model species. Yet, limited information is available on how closely cells and molecules of the anterior segment in commonly used models resemble those of humans. To address this issue, we profiled cells dissociated from anterior segment tissues of 4 model species: the Rhesus macaque (*M. mulatta*, 5158 cells); the cynomolgus Macaque (*M. fascicularis*, 9155 cells); the common swine (*S. scrofa*, 6709 cells); and mouse (*M. musculus*, 5067 cells). These species are among the most commonly used for studies on glaucoma: rhesus and cynomolgus monkeys are frequently used for basic studies of primate visual physiology and preclinical tests, respectively (Picaud et al.,2019); porcine anterior segments are used in aqueous outflow studies (Bachmann et al., 2006); and the broadest range of genetically modified lines is available in mice (Fernandes et al., 2015). We used a machine learning algorithm (XGBoost; Chen and Guestrin, 2016; see Methods) to find correspondences of cell types in the model species with those in humans. While significant conservation was noted among certain cell types across species, there were also notable differences.

#### M. mulatta

Cells collected from anterior chamber angle structures of *M. mulatta* clustered into 15 types **(*Figure 6A*)**. We identified 3 TM-like clusters in this species, similar to humans **(*Figure 6B*)**. One cluster corresponded to human Beam Cell A (mmC4), sharing markers including *BMP5* and *EDN3*; another corresponded to human JCT (mmC10) sharing preferential expression of markers including *ANGPTL7* and *CHI3L1*. The third cluster (mmC1), which we call Beam Cell X, could not be mapped to a single human cluster, but instead shared multiple markers of all human TM types; it was distinguished from mmC4 and mmC10 by its preferential expression of *CYP1B1* and *MGARP*. We identified types mapping 1:1 to human Schlemm canal (mmC7), vascular endothelium (mmC8), and the uveoscleral outflow pathway including ciliary muscle (mmC2), melanocytes (mmC9), myelinating Schwann cells (mmC5) and non-myelinating Schwann cells (mmC14) (***Figure 6F-K***). Two types of pericytes were present (mmC11, mmC12) compared to one in humans. Among immune cells, macrophage and NK/T clusters were present. B cells, mast cells and neurons were not detected, possibly due to their sparsity. Finally, due to targeted dissection of the TM strip in this species, collector channel endothelium, corneal epithelium and Schwalbe line cells were not recovered.

**Figure 6.**
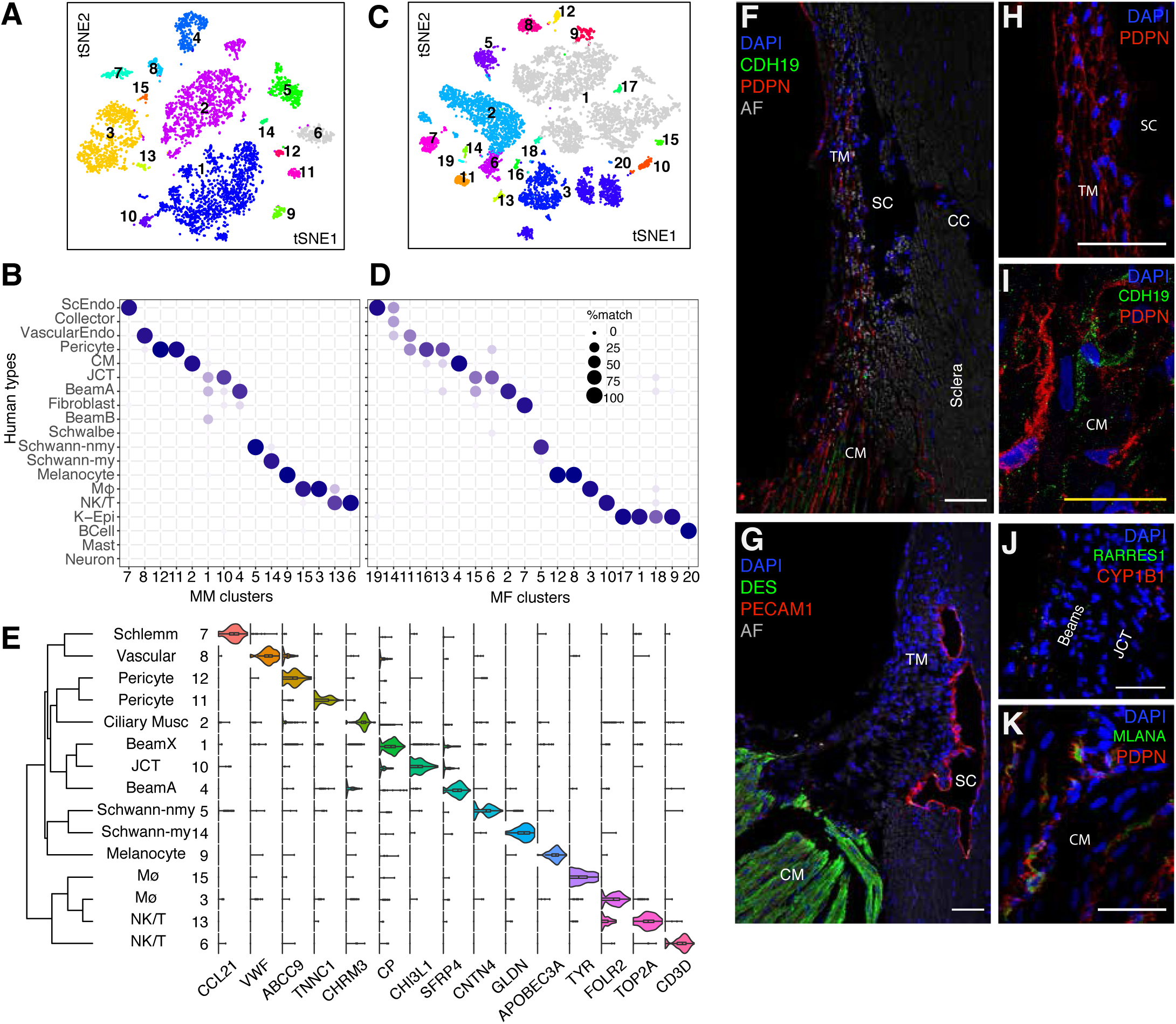
Cell types and gene expression by cells of the outflow pathways in 2 Macaque Species (*Mulatta* and *Fascicularis*) **A** tSNE plot showing 15 cell types derived from TM and associated structures of *M. mulatta* **B** tSNE plot showing 19 cell types derived from TM and associated structures of *M. fascicularis* **C** Transcriptional correspondence between human and *M. mulatta* cell types, summarized as a ‘‘confusion matrix.’’ In this and subsequent figures, the size of the circle and its intensity indicate the percentage of cells of a given cluster from the model species (column) assigned to a corresponding human cluster (row) by a classification algorithm trained on the human cells. **D** Transcriptional correspondence between human and *M. fascicularis*, shown as in **C**. **E** Violin plot showing examples of genes selectively expressed by each cell type in *M. mulatta*. **F-K** Histological localization of cluster markers in meridional sections of iridocorneal angles from *M. fascicularis* confirms identities of computationally derived cell types. *PDPN* stains trabecular beams (**F, H**) but also melanocytes within the ciliary muscle (CM) identified through co-staining with *MLANA* (**K**); Schlemm canal is outlined with *PECAM1*+ immunostaining whereas CM is highlighted with immunostaining against *DES* (**G**); CDH19 highlights Schwann cells in CM (**F,I**). Fluorescent RNA *in situ* hybridization against *RARRES1* (green) and *CYP1B1* (red) highlights cells in the TM (Beams > JCT). Bars show 50µm in F-H, J and K, and 25µm in I.

#### M. fascicularis

Cells collected from anterior chamber angle structures of *M. fascicularis* clustered into 19 types ***(Figure 6C)***. We identified 3 trabecular meshwork-like clusters (mfC2, mfC6, mfC15). Of these, one cluster, mfC2, preferentially expressed *BMP5* and *EDN3* and could be confidently mapped to human Beam Cell A (***Figure 6D***). The other two clusters (mfC6 and mfC15) expressed genes observed in multiple human TM clusters including *MYOC, MGP* and *ANGPTL7;* mfC6 tended to express more genes overlapping with the human JCT cluster (e.g. *FMOD, CEMIP*) and mfC15 a combination of human beam cell, fibroblast and Schwalbe line markers (e.g. *ANGPTL5, COL6A3, AQP1*, and *POU3F3*) ***(Figure S2A)***. Three vascular endothelial clusters were identified (mfC11, mfC14, mf19); these corresponded to the human capillary endothelium, collector channels and Schlemm canal endothelium, respectively. Other clusters mapped closely to cell types in the human dataset, including ciliary muscle (mfC4), Schwann cells (mfC5), macrophages (mfC3), B cells (mfC20) and NK/T cells (mfC10), melanocytes (mfC8, mfC12) and pericytes (mfC13, mfC16). No neurons were identified.

Based on results from human, we profiled cells from one sample that included the residual corneoscleral rim in addition to TM strips in hopes of identifying collector channel endothelium and matrix fibroblasts. Indeed, both additional cell types were recovered, in addition to multiple epithelial cell types (not described here). Matrix fibroblasts appeared during the initial unsupervised clustering as mfC7. Collector channel endothelium (mf14) could be identified within the original Schlemm canal cluster using a supervised approach based on differential expression patterns identified in humans ***(Figure S2B)***.

#### S. scrofa

Molecular profiling of porcine anterior segment tissue yielded 15 clusters ***(Figure 7A)***. Two clusters comprised TM cells. One mapped to human beam cell A (ssC1), with differential expression of *SFRP4* and *TMEFF2* in addition to *PON1, FMO1 and RARRES1*. The other mapped to human JCT (ssC3), selectively expressing *CCN3, FMO3, CEMIP* and *BMP3* ***(Figure 7B,C)***. While expression of *MYOC* was low across all clusters, it was preferentially expressed in ssC1. Both TM clusters expressed *CYP1B1, MGP, CFH*, and *NR2F1*.

**Figure 7.**
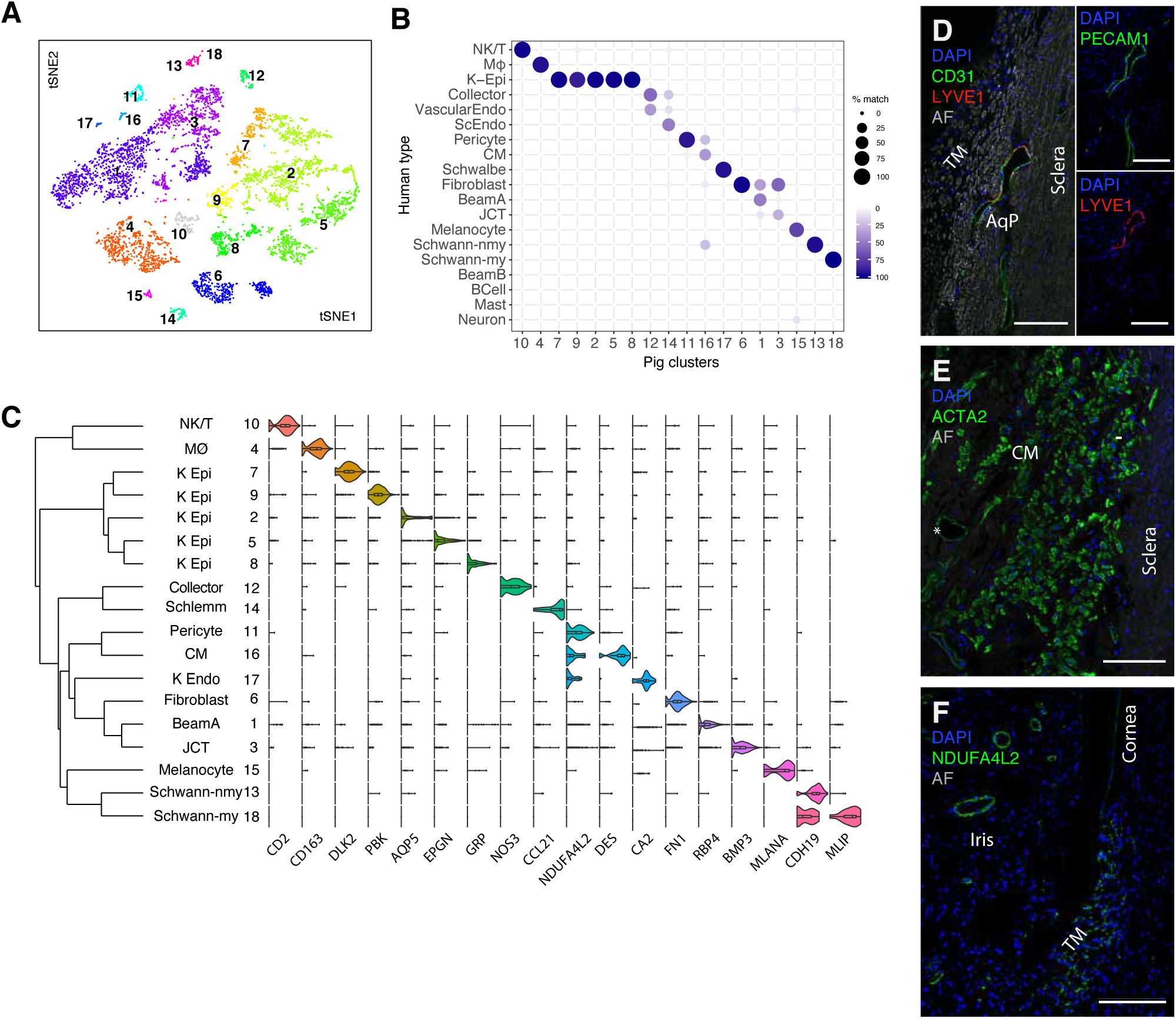
Cell types and gene expression by cells of the outflow pathways in the pig. **A** tSNE plot showing 18 cell types derived from TM and associated structures of pig. **B** Transcriptional correspondence between human and pig cell types, shown as in Fig. 6C. **C** Violin plot showing examples of genes selectively expressed by each cell type in pig **D** Immunostaining against *PECAM1* (green) highlights both the Aqueous plexus (AqP) and a downstream collector channel/scleral vessel whereas *LYVE1* stains only the AqP. **E** Immunostaining against *ACTA2* (green) highlights ciliary muscle and pericytes (asterisk). **F**. Immunostaining against *NDUFA4L2* (green) is strongest at the border of iris vessels, indicating pericytes, but also shows weak staining in the TM and corneal endothelium, consistent with transcriptional data. Bars show 50µm

Of 2 endothelial clusters identified, one mapped to human Schlemm Canal (ssC14) and the other to capillary endothelium (ssC12). While the pig does not possess a continuous SC and instead has more discontinuous vessels referred to as an aqueous plexus, cells lining this structure resemble those of human SC. The SC was distinguished from capillary endothelium through differential expression of lymphatic markers *CCL21, PROX1* and *LYVE1*, and was histologically validated with immunostaining against *LYVE1* (***Figure 7D***).

A ciliary muscle cluster (ssC16) was marked by preferential expression of DES and ACTA2, both of which we confirmed with immunostaining; the latter also stained pericytes (***Figure 7E***). *NDUFA4L2* was expressed predominantly by pericytes (ssC11) and CM but also corneal endothelium and to a lesser degree TM cells (***Figure 7F***). The low yield of ciliary muscle in our dissection can be attributed to the porcine angle anatomy, which differs from that of primates in having no clear anatomical connection between the ciliary muscle and conventional drainage structures (McMenamin and Steptoe, 1991). It is therefore easily excluded during dissection.

Other clusters corresponding to human cell types included myelinating (ssC18) and non-myelinating (ssC13) Schwann cells, melanocytes (ssC15), macrophages (ssC4) and NK/T cells (ssC10). Five clusters were derived from ocular surface epithelium due to a permissive dissection technique in which the TM/corneoscleral rim was dissociated. Cell types absent in this collection included Mast cells, B cells and neuronal types.

#### M. musculus

The mouse eye is small compared to those of other species profiled, so we dissociated the entire anterior segment, including cornea, iris and ciliary body. Presumably for this reason, initial clustering by cell class signatures revealed that 8766 of 13833 single cell profiles corresponded cells of the ocular surface epithelium. We eliminated these cells and re-analyzed the remaining 5067 cells, yielding 20 clusters **(*Figure 8A*)**.

**Figure 8.**
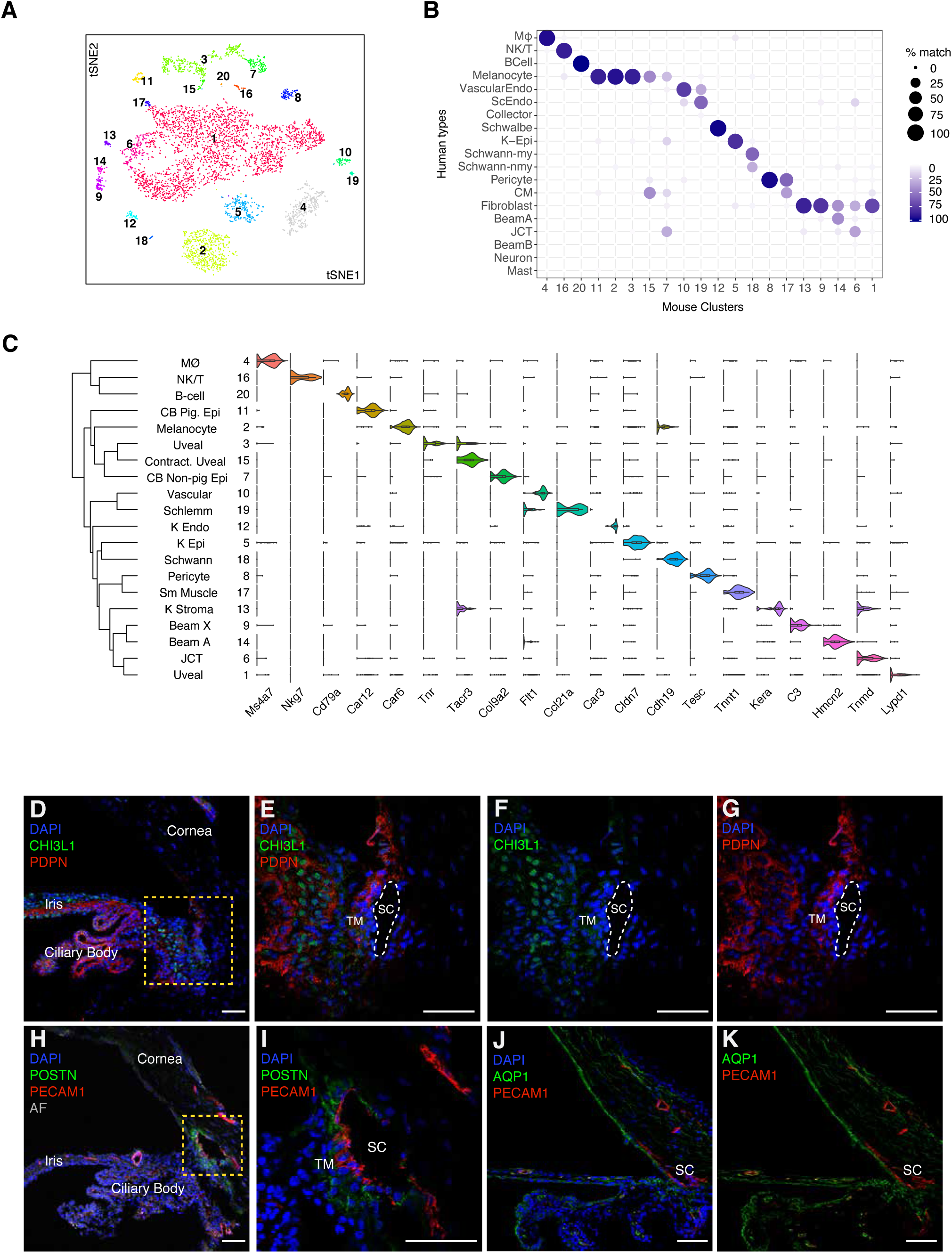
Cell types and gene expression by cells of the outflow pathways in the mouse. **A** tSNE plot showing 20 cell types derived from TM and associated structure of mouse. **B** Transcriptional correspondence between human and mouse cell types, shown as in Fig. 6C. **C** Violin plot showing examples of genes selectively expressed by each cell type in mouse. **D-G** Pdpn is present in multiple cell types including pigmented and nonpigmented epithelium of the iris and CB as well as a subset of TM cells, K Endo, and K Epi, whereas *Chil1* (ortholog to *CHI3L1* in humans) stains a different subset of TM cells and to a lesser extent cells within the CB. **E-G** Higher magnification of boxed area. **H-I**. Immunostaining against Postn (green), a secreted protein, highlights Schlemm canal and JCT cells. Pecam1 (red) highlights SC as well as vascular endothelial clusters. J-K Immunostaining against Aqp1 highlights corneal clusters (K stroma, K Endo) most vividly, in addition to iris, ciliary body and TM cells. Bars show 50µm

Although the mouse iridocorneal angle is more compact than that of humans, anatomical studies suggest that conventional outflow structures in mice are similar in many respects, with trabecular lamellae, distinctive juxtacanalicular tissue, and a continuous Schlemm canal (Overby et al., 2014). We identified three candidate TM clusters (mC6, mC9, and mC14) through their high levels of *Myoc* expression. Two of these clusters, mC14 and mC6, mapped preferentially to human Beam cell A and JCT, respectively, while also demonstrating some ECM expression patterns similar to the human corneoscleral fibroblast cluster **(*Figure 8B*)**. The third candidate TM cluster, mC9, despite sharing many markers with fibroblasts (e.g. *Pi16, Fbn1, Mfap5, Tnxb, Clec3b*), was closely related to mC14 and thus tentatively named Beam Cell X. All three TM clusters also expressed *Mgp* and *Pdpn* similar to human TM cells; the latter was confirmed histologically with immunostaining, as was expression of *Chil1* (*CHI3L1*) in mC9 and mC14 ***(Figure 8D-G, Figure S4)***. Other DE genes within the mouse JCT cluster included *Nell2, Chad* and *Tnmd*, whereas DE genes within the mouse Beam clusters included *Sfrp4* (both mC9 and mC14), *Tmeff2* (mC14) and *Fmo2* (mC14).

Among other outflow cell types, we identified 2 separate vessel clusters (mmC10, mmC19) corresponding to vascular and Schlemm canal endothelium, respectively. *Postn*, which was also differentially expressed within the human Schlemm canal cluster, was also expressed in mouse Schlemm canal as well as JCT **(*Figure 8H,I*)**.

Other clusters mapping to human cell types included corneal endothelium / Schwalbe line (mC12), smooth muscle (mC17), pericytes (mC8), Schwann cells (mC18) and uveal melanocytes (mC2). (**Figure 8J,K**) In contrast to our human data, the smooth muscle and Schwann cell clusters were small, which can be attributed in large part to the diminutive murine accommodative apparatus (Overby et al., 2014).

Owing to the dissection method, the mouse dataset also included a large number of cells derived from iris and ciliary body. We tentatively identified these clusters based on markers identified in previous studies (Diehn et al., 2005, Janssen et al., 2012); they comprised melanocytes, stromal fibroblasts, pigmented epithelium and nonpigmented epithelium of the iris and ciliary body.

### Conservation of expression patterns among species

Next, we assessed expression patterns in model species of key genes selectively expressed in cell types of the human aqueous humor outflow pathways. In general, conservation was striking.

Many markers expressed across all human trabecular meshwork clusters **(*Figure 9A*)** were conserved in other species **(*Figure 9A*)**. They included matrix-related genes such as *DCN* and *MGP* and the retinoic acid-related genes *RARRES1* and *RBP*. However, some differences among species were evident. *PDPN*, a selective marker for TM cells in humans, was less selective for these cells in other species. For example, it was also found at similar levels in uveal melanocytes in all four non-human species. *BMP5*, expressed in Beam Cell A in human and macaque, was present but less specific in pig and absent in mouse. The human JCT marker, *CHI3L1*, was preferentially expressed in JCT clusters in monkey and mouse but was notably absent in pig.

**Figure 9.**
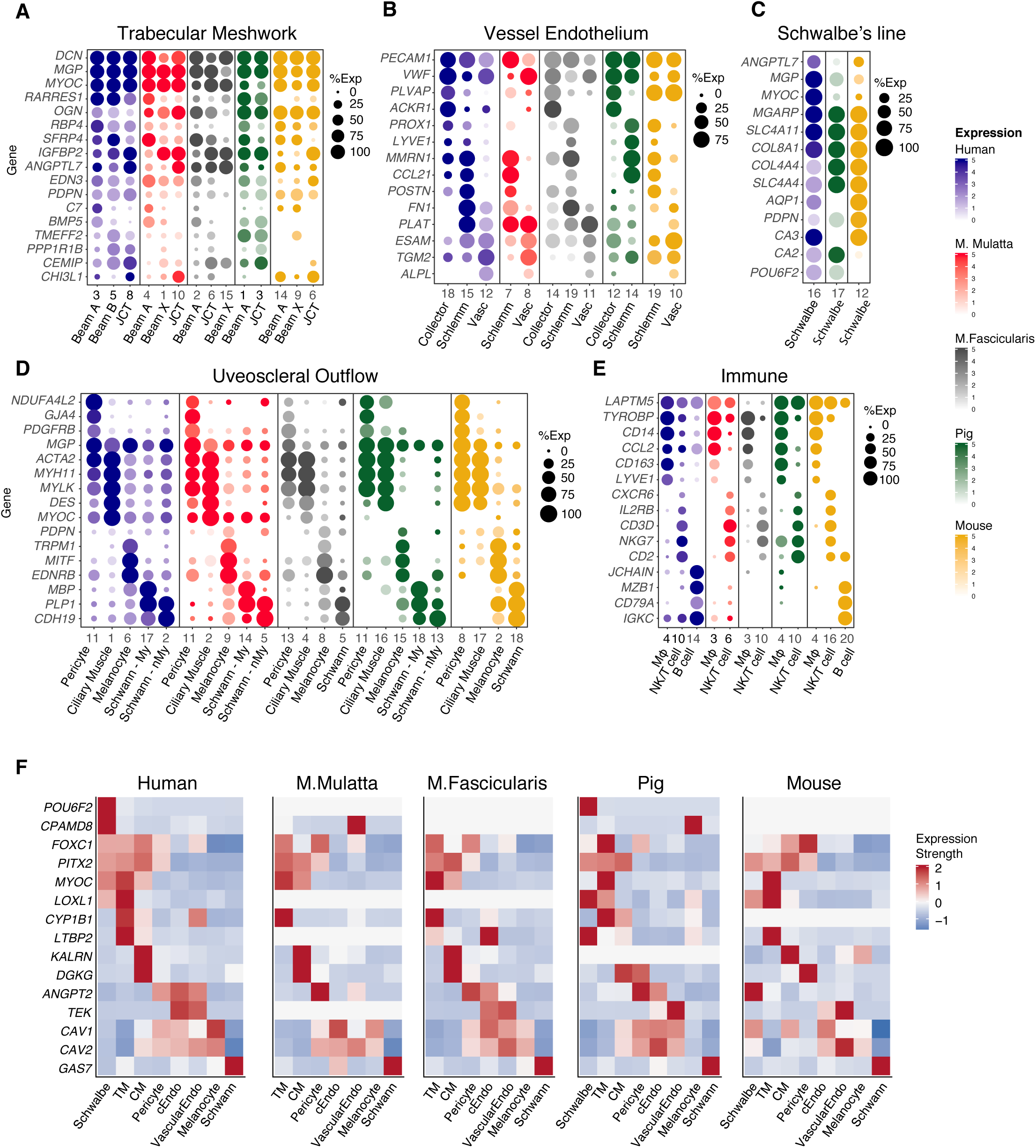
Comparison of gene expression across species. **A-E** Key genes are shown in dot plots for cell types comprising the trabecular meshwork (A), vascular endothelium (B), Schwalbe line / Corneal Endothelium (C) and uveoscleral outflow pathway (D) as well as immune cells (E). **F** Heat map showing expression of genes implicated in human POAG in aqueous outflow cells of human (replotted from Fig S1) and model species.

There was also excellent correspondence between markers of for other cell types that comprise the conventional pathway. Markers of collector channel, Schlemm canal and vascular endothelium markers were generally well conserved among humans and macaque species (***Figure 9B***). Pig and mouse also shared many similar markers; however, Schlemm canal in these species tended to demonstrate more prominent expression of lymphatic markers than primates. Among species in which Schwalbe line / corneal endothelial cells were obtained, there was also good correspondence **(*Figure 9C*)**. Of note, whereas the Schwalbe line cluster in humans includes cells at the junction between cornea and TM, the corresponding clusters in pig and mouse more likely represent a larger proportion of corneal endothelia. Markers of cell types comprising the uveoscleral pathway and immune cells were also very well conserved among species (***Figure 9D,E***).

Finally, we analyzed patterns of disease gene expression could be identified in these model species ***(Figure 9F)***. To facilitate comparison, individual TM cell clusters from each species were merged during this analysis, and a threshold was set such that only genes with expression in >10% of cells in a minimum of one cluster were plotted. While some genes were consistently expressed in corresponding cell types across species, others exhibited significant differences. For example, we noted that the classic anterior segment dysgenesis genes *FOXC1* and *PITX2* were reliably expressed in trabecular meshwork cell types across all species. Similarly, *MYOC* was also expressed in TM cells across species; however, in this case, subtle differences in other cell types were observed. For example, in human and macaque, *MYOC* was strongly expressed in both TM and ciliary muscle, whereas in pig and mouse it was expressed only in TM. Similarly, *MYOC* was highly expressed in Schwalbe line cells of human but not in the corresponding corneal endothelial clusters of pig and mouse (as mentioned above, likely because these represent a larger set of corneal endothelia, not just the peripheral subset). *LOXL1*, while expressed in TM clusters of human, pig and mouse, was absent in the TM of both Macaque species. *KALRN*, expressed in human, macaque and mouse CM, was absent in pig. Finally, *CYP1B1*, expressed in TM cells across human, macaque and pig, did not meet threshold expression levels in mouse TM. This is consistent with prior studies, which have reported *CYP1B1* cDNA but not protein expression in human adult TM samples, and negative IHC staining in mouse TM (Vasilou and Gonzalez, 2008).

## DISCUSSION

We used scRNAseq to profile cells comprising the aqueous humor outflow pathway in humans, generating a cell atlas for these tissues and identifying new markers for each cell type. We then used the atlas to localize the expression of genes implicated in glaucoma. These findings offer new insights into molecular architecture of the TM cells and highlight potential roles in IOP homeostasis for non-TM cell types in the anterior chamber angle. Finally, we profiled cells in the outflow pathways of four model species – *M. mulatta, M. fascicularis, S. scrofa, and M. musculus* – providing a foundation for using these models in studies on regulation and dysregulation of IOP.

### Technical issues

A major challenge in human vision research is that essentially all ocular tissues must be obtained either post-mortem or post-enucleation. Furthermore, post-mortem ocular tissues of the anterior segment suitable for transplantation are understandably prioritized for this purpose over research. In this study, all post-mortem tissues sequenced were obtained within ∼6 hours of death from a Rapid Autopsy Program enrolling predominantly oncology patients. While donors had no documented pathological ocular history or clinical evidence of eye disease on examination, they did have varying degrees of chronic systemic disease and different levels of antemortem exposure to chemotherapeutic and steroid medications and this represents a limitation of our study. Furthermore, lack of documented or physically evident ocular pathology cannot be taken as evidence that none existed. Since we focused primarily on cell type classification in this study, rather than quantitative determination of gene expression levels, we believe that the donors’ systemic diseases did not influence the ultimate cell atlas. This was further corroborated in two ways: (1) demonstrating that the same cell types were obtained from all individuals and (2) performing histological validation on tissues obtained from a wider variety of donors through an eye bank, as well as on separate rapid autopsy patients.

### Human cell atlas

In the human AH outflow pathway, we identified 19 major cell types: 8 comprising the conventional outflow pathway (including 3 distinct populations within the filtering trabecular meshwork), 7 comprising the uveoscleral pathway, and 4 immune cell populations.

By histological criteria, the filtering trabecular meshwork has been divided into three layers: a uveal layer adjacent to the anterior chamber, a juxtacanalicular layer adjacent to Schlemm canal, and a corneoscleral layer in between (Tamm, 2009; Stamer and Clark, 2017). The JCT cells in our dataset clearly localized closest to Schlemm canal, with the Beam A and B cells localizing to the other two layers. It is tempting to assign Beam A and B to uveal and corneoscleral layers, respectively, but our histological analysis suggests that they are in fact intermingled.

The lymphatic marker podoplanin (*PDPN*, aka *D2-40*) emerged as a robust marker for all human TM cell types (clusters 3, 5, and 8) consistent with previous studies (Birke et al., 2010, Watanabe et al., 2010). Along with *CCL2* (a chemoattractant for monocytes) and *VCAM1* (a mediator of immune cell migration), there was a marked specificity of *PDPN* expression among cells of the conventional pathway as compared to the uveoscleral pathway, suggesting that the former acts as an immunological “sink” guiding antigen presenting cells and other immune cells toward SC, the venous system, and ultimately toward the spleen (Aspelund et al., 2014). Consistent with this idea, Schlemm canal expresses markers of lymphatic vessels (e.g., *CCL21* and *FLT4/VEGFR3*, whereas collector channels express venular markers (e.g. *ACKR1*).

### Model species

In comparing the cellular composition of human AH outflow pathways to those of commonly used models, we documented excellent correspondence for many cell types across species. Ciliary muscle cells, pericytes, melanocytes, and Schwann cells were present in all species and demonstrated shared expression of canonical markers. Corneal endothelium was identified in pig and mouse, but in neither macaque species, likely due to our targeted dissection technique. Two or more types of vascular endothelium, defined as PECAM1+ TIE1+ clusters, were also present in each species, and could be assigned either to Schlemm canal or other vasculature. Among immune cells, Lyve1+ CD163+ CD68+ macrophages were also present in all species; other immune cells were identified in each species with less correlation, perhaps due to their relatively sparse numbers. Similarly, neurons were limited to the human dataset most likely due to their low numbers.

On the other hand, while trabecular meshwork cells could be identified in each species, they demonstrated substantial variability across species. Three TM types –two beam cell types and one JCT cell type – were present in all three primates, but they failed to map 1:1 across species. Beam A and JCT cells were present in all three species, but the second beam type, which we call Beam X, appeared to be intermediate in expression pattern between human Beam A and Beam B. Moreover, we identified only one beam cell type in pig and mouse, although it is possible that further analysis with increased sample size would allow a subdivision of this cluster. Some DE markers, such as *PDPN, RARRES1, CHI3L1*, and *ANGPTL7*, which were selectively expressed by beam and/or JCT cells in human tissue, were either absent in the model species, or nonspecific. Others were conserved across species, including *MYOC, EDN3*, and *RBP4*. Interestingly, *PDPN* did not demonstrate the same specificity in any of our model species, suggesting that this may be a human-specific feature.

Genes involved in the retinoic acid pathway were expressed in the conventional outflow tract across species. They included *RARRES1, RARRES2, RBP4, FABP5, ADH1B* and the glaucoma-associated disease gene CYP1B1 **(Figure S2K,L)**, which converts retinol to retinoic acid (Chambers et al., 2007). In addition to a possible role for retinoic acid in maintaining a tolerogenic microenvironment within the anterior chamber, retinoic acid response genes have been implicated as potentially important mediators in the steroid-induced upregulation of *MYOC* (Prat et al., 2017).

### Glaucoma

Glaucoma is a phenotypically heterogeneous disease with traits dictated by complex interactions among age, environment and genes. Examination of cell-type specific expression patterns of both Mendelian genes and GWAS susceptibility loci revealed multiple expression patterns, of which we note four groups. First, several IOP-associated disease genes were selectively expressed by TM cell types (e.g., *CYP1B1* and *EFEMP1*), supporting their potential contribution to function and or/dysfunction of this tissue. Second, others were expressed not only by TM but also cells of the uveoscleral pathway (e.g., *MYOC, PITX2*, and *FOXC1*). Third, a few genes implicated in high IOP mapped selectively to cells of the uveoscleral pathway (e.g., KALRN). Together, these results may indicate an integral contribution of uveoscleral pathway to disorders of IOP. Finally, some genes were expressed by RGCs, often in addition to TM cells. For example, *TMCO1*, while clearly expressed in the anterior segment outflow pathways, also demonstrated robust expression in RGCs, consistent with reports suggesting it to be linked more closely with inherent RGC susceptibility independent of IOP (Scheetz et al., 2016).

Although TM cells have been shown to exhibit phagocytic abilities, it is possible that resident macrophages in the conventional outflow pathway also contribute to the phagocytic workload at this site of filtration and that their stimulation or recruitment to the TM may serve lower IOP, as has been suggested to occur after Selective Laser Trabeculoplasty (SLT) (Alvarado et al., 2010). Similarly, dysfunctional macrophages may contribute to elevated IOP in some cases of secondary open-angle glaucoma (Hamanaka et al., 2002).

## MATERIALS & METHODS

### Tissue Acquisition

Human ocular tissues used for sequencing, immunohistochemistry and *in situ* hybridization were obtained from Massachusetts General Hospital in collaboration with the Rapid Autopsy Program, Susan Eid Tumor Heterogeneity Initiative. Eyes were collected a median of 6 hours postmortem (range 3-14hrs; see **Table S1**). The whole globe was immediately transported to the lab in a humid chamber on ice. Hemisection was performed at the pars plana and the anterior segment was then placed in Ames medium equilibrated with 95% O_2_/5% CO_2_. For sequencing, the following dissection was performed: after isolation of the anterior segment, the lens, iris, and ciliary body were removed with a gentle peeling method. The corneoscleral button was hemisected and a small wedge was set aside for fixation. Approximately 9 clock hours (or 270 degrees) of TM tissue was peeled from the scleral sulcus of the remaining tissue with jewelers’ forceps. A permissive dissection technique allowed for adjacent tissue from the ciliary muscle to be incorporated into the collection tube if it was liberated in conjunction with the TM strip.

Other human corneoscleral buttons used for IHC and *in situ* hybridization were provided by the Lions Vision Gift (Portland, OR) and were collected <16 hr postmortem and fixed in ice-cold 4% PFA. No ocular disease was reported in any of the human donors and no abnormalities were noted during microdissection. Donor details are provided in Table S1.

Non-human primate eyes were obtained from macaques 4 to 10 years of age that had reached the end of unrelated studies at supplying institutions. No ocular or visual abnormalities were noted. Data presented in this manuscript did not covary with any treatment that had been applied to the animals. For sequencing, two eyes from one female crab-eating macaque (*Macaca fascicularis*, 6 years of age) were used, and two eyes from one male rhesus macaque (*Macaca mulatta*). Eyes were collected either pre-mortem under deep anesthesia or post-mortem (≤ 45 min), after which a rapid hemisection was performed and the anterior immediately placed in ice-cold Ames solution (Sigma-Aldrich; equilibrated with 95% O_2_/5% CO_2_ for all use), where they were stored before experimentation. Experiments described below commenced within 6 hours of death.

Porcine (*sus scrofa*) eyes were obtained from a local abattoir, transported back to the lab in a humid chamber on ice, and dissected as above. For sequencing, 2 eyes from 2 individual pigs were used. Due to the porcine iridocorneal anatomy, TM strips were not dissected, and instead, corneoscleral wedges were trimmed and digested *in toto*.

Mouse eyes were collected from male and female 12-week-old CD1 mice obtained from Charles River Laboratories. After euthanasia, eyes were enucleated and transferred to Ames medium equilibrated with 95% O_2_/5% CO_2_. The anterior segment was dissected from the posterior portion of the eye with microscissors, and the lens was gently removed with forceps. The entire anterior segment, including cornea, iris, ciliary body, and the TM was digested.

### Single cell isolation

Tissues were digested enzymatically for 30 minutes at 37°C with papain (Worthington, LS003126) 20 units/mL in Ames. Following digestion, the tissues were triturated into single cell suspensions with 0.15% ovomucoid and 0.04% bovine serum albumin (BSA) in Ames solution and filtered through a 40 um strainer. Single cell suspensions were diluted at a concentration of 500-1800 cells/µL in 0.04% non-acetylated BSA/Ames for loading into 10X Chromium Single Cell Chips. (Zheng et al., 2017). Data were obtained using both V2 and V3 kits and specified in Table S1.

### Droplet-based scRNA-seq

Single cell libraries were generated with either Chromium 3’ v2 or V3 platform (10X Genomics, Pleasanton, CA) following the manufacturer’s protocol. Briefly, single cells were partitioned into Gel bead in Emulsion (GEMs) in the GemCode instrument with cell lysis and barcoded reverse transcription of RNA, followed by amplification, shearing and 5’ adaptor and sample index attachment. On average, approximately 10,000 single cells were loaded on each channel and approximately 6,000 cells were recovered. Libraries were sequenced on Illumina HiSeq 2500.

### Histology

Corneoscleral wedges were fixed in 4% PFA for 2-24 hr, transferred to PBS, sunk in 30% sucrose overnight, then embedded in tissue freezing medium and mounted onto poly-d-lysine coated slides in 20 μm meridional sections with ProLong Gold Antifade (Invitrogen). For IHC, slides were incubated for 1 hr in protein block, overnight with primary antibodies, and 2 hr with secondary antibodies. Initial block and secondary antibody incubation were done at room temperature and primary antibody incubation at 4°C. Single molecule fluorescent *in situ* hybridization was performed using commercially available RNAScope Multiplex Fluorescent Assay V2 (Advanced Cell Diagnostics, Newark, CA). Briefly, slides were baked at 60°C for 30 minutes and incubated for 10 minutes at RT with hydrogen peroxide. Two Protease III incubations were performed (30min, then 15 min). Probe hybridization and subsequent steps were per standard manufacturer protocol. Antibodies and *in situ* probes are catalogued in **Table S2**.

### Image acquisition, processing and analysis

Images were acquired on Zeiss LSM 710 confocal microscopes with 405, 488-515, 568, and 647 nm lasers, processed using Zeiss ZEN software suites, and analyzed using ImageJ (NIH). Images were acquired with 20X, 40X or 100X oil lens at the resolution of 1024×1024 pixels, a step size of 1.0 μm, and 90μm pinhole size.

### Computational Methods

#### Clustering analysis

Sequencing data was demultiplexed and aligned using the cellranger (10X Genomics) mkfastq and count functions respectively (cellranger version 2 for samples collected with v2 kit and version 3 for samples collected with v3 kit). Reads were aligned to the following reference genome: Human samples-GRCh38, Macaca Mulatta-Mmul8, Macaca Fascicularis-MacFas5 with our augmented transcriptome file (Peng *et al*, 2019), Pig-Sscrofa11, Mouse-mm10. The number of genes/transcripts detected per cell was plotted for every sample and a threshold was chosen based on their distributions for each species. Cells with fewer than 600 genes were excluded from further analysis for primates and pig; the threshold for mouse was 1000 genes/cell.

The analysis pipeline follows methods described by Peng *et al*. (2019). (1) For each species, the UMI count matrix was normalized by the total number of UMI counts for each cell and multiplied by a scale factor, the median UMI counts of the group to correct library size differences. The normalized UMI count matrix was then log transformed after adding 1. (2) High Variable Genes (HVG) were identified using method described in Pandey *et al*., 2018. For every gene, the mean (μ) and coefficient of variation (CV) of UMI counts among cells within each group was calculated. The excess CV (eCV) was measured by subtracting the predicted value from a Poisson-Gamma mixture based null model of CV v.s. μ. HVGs were defined as genes whose eCV> [mean of eCV] + 1.3*[SD of eCV]. (3) Batch correction was performed on the expression matrix of HVGs using linear regression model adapted from the ‘RegressOut’ function in the ‘Seurat’ R package. (4) Principal Component analysis (PCA) was performed on the batch-corrected expression matrix of the HVGs. The top 90 PCs were calculated using the ‘irlba’ R package on centered and scaled data. Statistically significant PCs were estimated based on the Tracy-Widom distribution (Patterson et. al., 2006). The PCA-reduced data was projected to 2D using t-distributed stochastic neighbor embedding (t-SNE) for visualization. (5) Unsupervised Louvain-Jaccard clustering method was applied to cells in the PCA reduced dimensional space of the significant PCs (Shekhar et al, 2016). (6) Clusters formed by either low quality cells or doublets were removed before further analysis. The former could be identified with lower median number of genes/transcripts per cell, higher proportion of reads from mitochondrial genes, and lack of uniquely expressed genes. The latter could be identified as overall higher number of genes/transcripts than other clusters, expression of marker genes from multiple types, and also lack of uniquely expressed genes. (7) To assess the overall relationships among clusters, a dendrogram was built in each species using normalized expression matrix of HVGs. (8) To correct for potential over-clustering due to batch effect, or the lack of computation power in distinguishing types, the following steps were taken. To address the first issue, cluster pairs that mapped closest to each other on the dendrogram were tested, and they were merged iteratively until sufficient differences were found between the current cluster pair (≥5 DE genes enriched both ways, log fold-change >1.2, adjusted p value < 0.001; statistical testing was performed using the ‘MAST’ R package). After each merge, a new dendrogram was built for testing the new pair. To address the second issue, sub-clustering was performed on each cluster with newly defined HVGs among cells and using previous described testing method for verification. In most cases, result from sub-clustering represented over-clustering; however, in some cases, we were able to distinguish real types (i.e. the human Schlemm canal and collector channel cells were grouped as one cluster at the initial clustering, but split into two at the sub-clustering step). In other cases, the lack of clustering power was due to limited sample size, as in all the species we analyzed except the human dataset. Referring to the cross species mapping (described below) to first spot the cluster as a potential cell type mixture in non-human dataset, we used supervised methods to separate them with type markers found in human. For example, the Schlemm canal and collector channel in *M. fascicularis* were initially clustered as a single type using unsupervised methods, but the human type markers were present in separate cell populations (FigureS2B), allowing us to split the cluster. Subsequently, criteria described above were applied to verify the new clusters. (9) Due to differences in the dissection method, we obtained a large amount of corneal epithelium from our mouse collection. The presence of these cells saturated the dynamic range in feature space and weakened our computational power in classification of other types; therefore the majority of corneal epithelium was removed at the initial analysis.

#### Cross species comparison

To evaluate the transcriptomic similarity between human and other model species, we applied a boosted-tree based machine learning algorithm XGBoost (Chen and Guestrin, 2016) using the ‘xgboost’ R package. We used the human dataset for training because it was the largest. Classifiers were trained on 80% of the human dataset using the HVGs shared between human and each of the other species. The remaining 20% was used as a validation dataset to evaluate the accuracy of the classifier; for all four classifiers built, the validation accuracy was >95%. Data from non-human species were used as the testing dataset whereby cells that get more than a 16% vote (calculated from the equation below) for the ‘winner’ type are assigned with human type identity, while those failing to pass the criteria are assigned as ‘unmapped’.

The criteria is calculated as promotion of winning votes ≥: 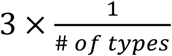, where # of types in the human dataset is 19. In some cases, a single cluster in the testing dataset might map to two types in the training dataset, and this serves as a clue of potential cell type mixture as mentioned previously. Further sub-clustering and verification was performed accordingly.

### Data availability

The accession number for the raw and unprocessed data files reported in this paper is GEO: GSExxxxxx.(in process). Data can be visualized at the Broad Institute’s Single Cell Portal at https://portals.broadinstitute.org/single_cell.

### Study Approval

All animal procedures performed in this study were conducted in compliance with the Association for Research in Vision and Ophthalmology’s Statement for the Use of Animals in Ophthalmic and Vision Research and guidelines for the care and use of animals and human subjects at Harvard University and Partners Healthcare. Acquisition and use of human tissue was approved by the Human Study Subject Committees (DFCI Protocol Number: 13-416 and MEE - NHSR Protocol Number 18-034H). Acquisition and use of non-human tissue was approved by the Institutional Animal Care and Use Committee (IACUC) at Harvard University.

## ACKNOWLEDGEMENTS

This work was supported by NIH (5K12EY016335-12 R21, EY028633 and U01 MH105960), the Chan-Zuckerberg Initiative (CZF-2019-002459) and the Klarman Cell Observatory of the Broad Institute of MIT and Harvard. We thank the donors and their families for their generosity. We are grateful to Michael Do for providing Macaque tissue. We thank Carl Romano and Dan Stamer for sharing results of their parallel study prior to submission.

