## Supplemental Figures and Tables for "Cell Atlas of Aqueous Humor Outflow Pathways in Eyes of Humans and Four Model Species Provides Insights into Glaucoma Pathogenesis"

#### SUPPLEMENTARY FIGURES AND TABLES

**Figure S1**
A. Beam cells immunostained with *PDPN* (green) and *FABP4* (red).
B. Enlargement of boxed area in A.
C. Ciliary muscle cells co-immunostained for CHRM3 and DES.
D. Pericytes immunostained for NDUFA4L2, wrapped around a vessel stained with PECAM1.
E. Neuron within the ciliary muscle immunostained for ELAVL4/HuD.
F. Mast cells within the TM immunostained for IL1RL1.
G. B cells within the TM immunostained for *CD27*.
H. Immunostaining against ADH1B highlights scleral fibroblasts in the vicinity of TM and SC. This image is shown in higher magnification in **3F**.
I. Fluorescent RNA *in situ* hybridization against *ANGPTL7* (green) and *CHI3L1* (red) highlights cells in the JCT. Schlemm canal outlined in dashed line. A cropped version of this image is shown in higher magnification in **3I**.
J. Fluorescent RNA *in situ* hybridization against *TMEFF2* (green) and *PPPR1B1* (red) highlights beam cells. A cropped version of this image is shown in higher magnification in **3H**.
K. Fluorescent RNA *in situ* hybridization against *RARRES1* (green) and *CYP1B1* (red) demonstrates positive signal within beam cells > JCT.
L. Enlargement of area boxed in K.

**Figure S2 Disease Genes**
Heat map showing expression of genes implicated in glaucoma by key cell types. Genes detected in more than 10% cells of at least one type were shown.

**Figure S3 Analysis of cell types and gene expression in M. Fascicularis**
**A** Violin plot showing examples of genes selectively expressed by each cell type in *M. fascicularis*
**B** Supervised clustering of Schlemm Canal and collector channels in *M. fascicularis*

**Figure S4 Expression of key marker genes in mouse**
Violin plots showing expression of key marker genes in mouse

**Table S1**
Information about donors used for RNA-seq. COD, cause of death.

**Table S2**
Information about donors used for histological validation.

### Figure S1

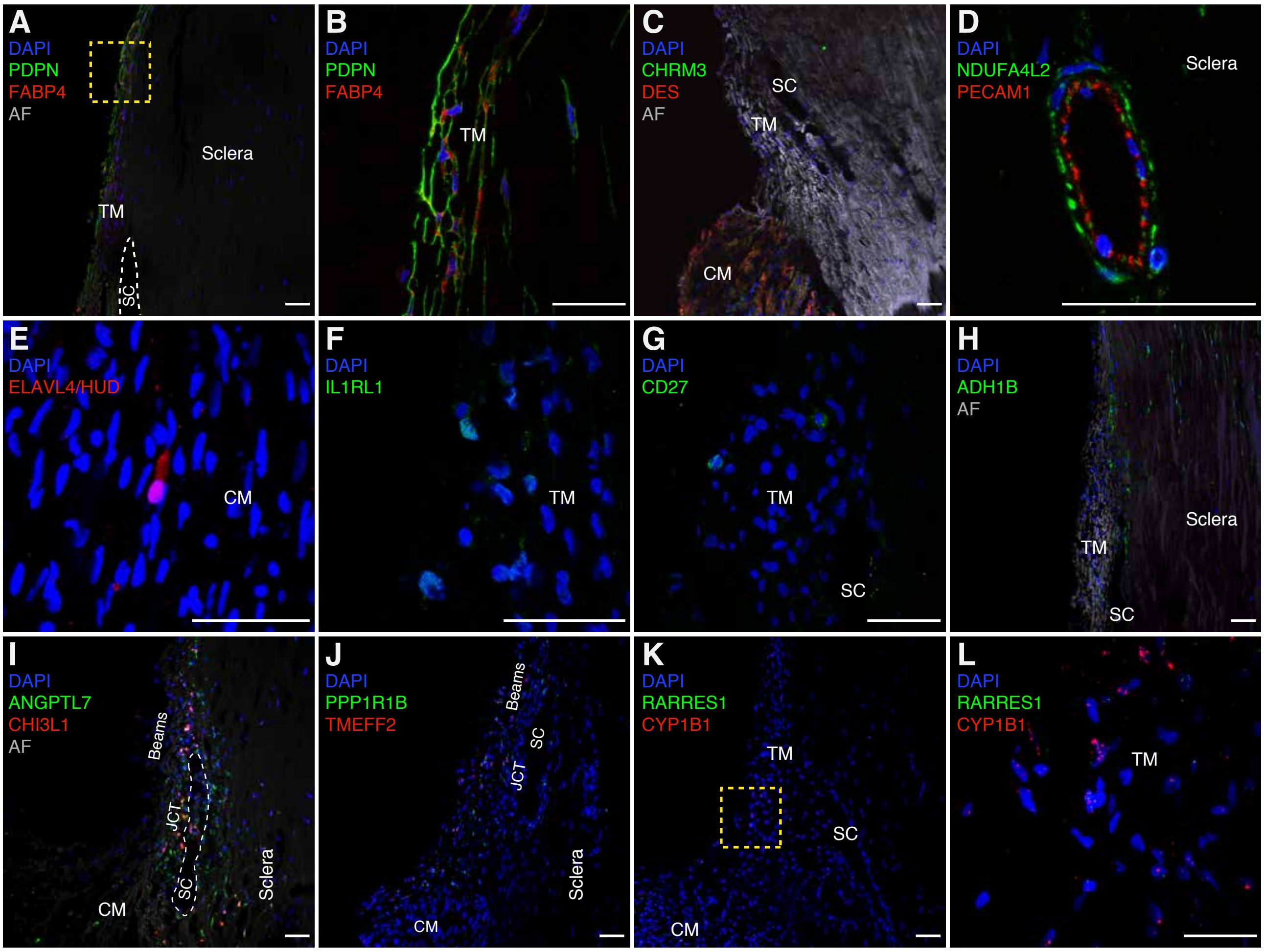
A
DAPI
PDPN
FABP4
AF
Sclera
TM
SC
B
DAPI
PDPN
FABP4
TM
C
DAPI
CHRM3
DES
AF
SC
TM
CM
D
DAPI
NDUFA4L2
PECAM1
Sclera
E
DAPI
ELAVL4/HUD
CM
F
DAPI
IL1RL1
TM
G
DAPI
CD27
TM
SC
H
DAPI
ADH1B
AF
Sclera
TM
SC
I
DAPI
ANGPTL7
CHI3L1
AF
Beams
JCT
SC
Sclera
CM
J
DAPI
PPP1R1B
TMEFF2
Beams
SC
JCT
Sclera
CM
K
DAPI
RARRES1
CYP1B1
TM
SC
CM
L
DAPI
RARRES1
CYP1B1
TM

### Figure S2

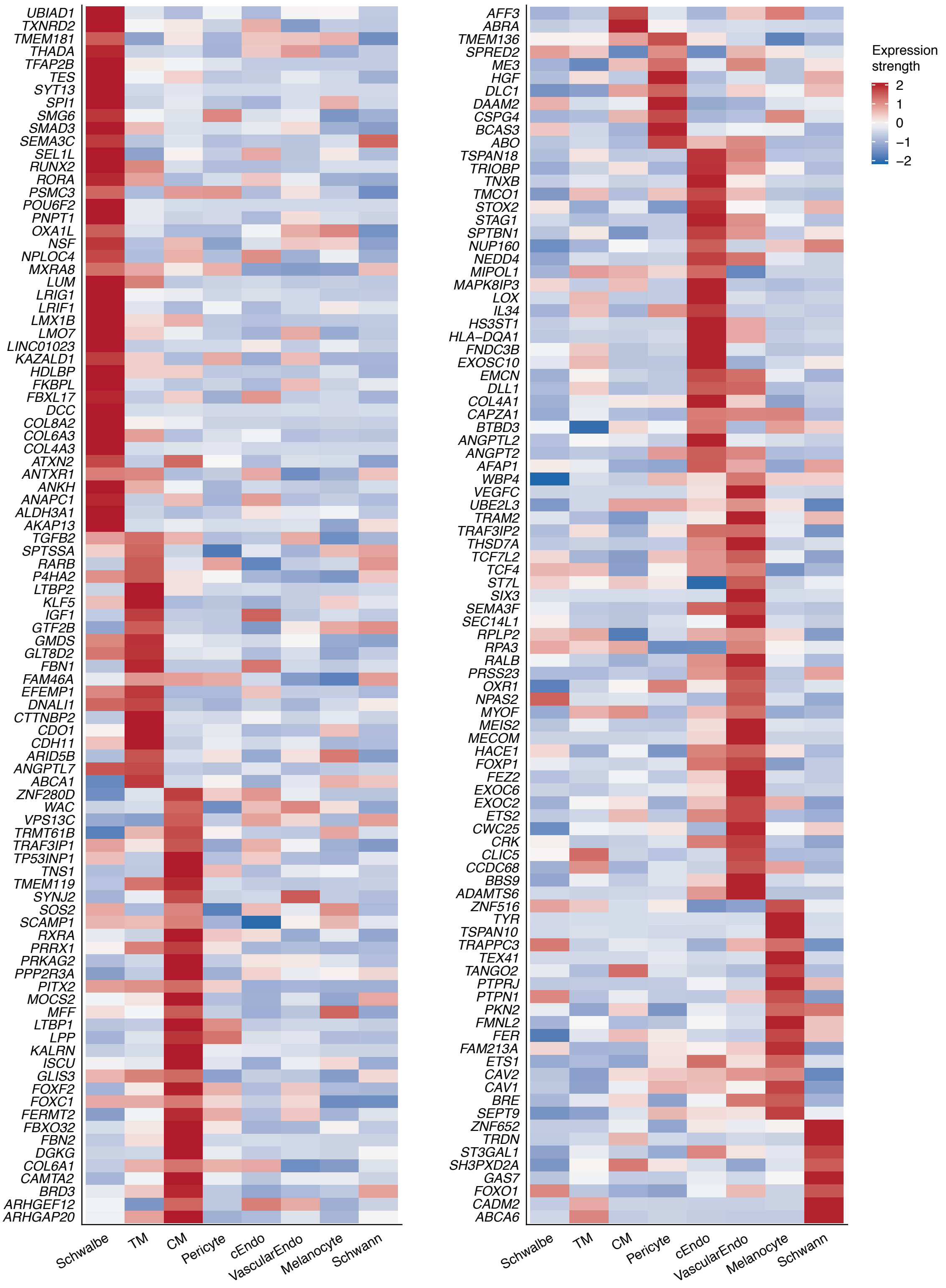
UBIAD1
TXNRD2
TMEM181
THADA
TFAP2B
TES
SYT13
SPI1
SMG6
SMAD3
SEMA3C
SEL1L
RUNX2
RORA
PSMC3
POU6F2
PNPT1
OXA1L
NSF
NPLOC4
MXRA8
LUM
LRIG1
LRIF1
LMX1B
LMO7
LINC01023
KAZALD1
HDLBP
FKBPL
FBXL17
DCC
COL8A2
COL6A3
COL4A3
ATXN2
ANTXR1
ANKH
ANAPC1
ALDH3A1
AKAP13
TGFB2
SPTSSA
RARB
P4HA2
LTBP2
KLF5
IGF1
GTF2B
GMDS
GLT8D2
FBN1
FAM46A
EFEMP1
DNALI1
CTTNBP2
CDO1
CDH11
ARID5B
ANGPTL7
ABCA1
ZNF280D
WAC
VPS13C
TRMT61B
TRAF3IP1
TP53INP1
TNS1
TMEM119
SYNJ2
SOS2
SCAMP1
RXRA
PRRX1
PRKAG2
PPP2R3A
PITX2
MOCS2
MFF
LTBP1
LPP
KALRN
ISCU
GLIS3
FOXF2
FOXC1
FERMT2
FBXO32
FBN2
DGKG
COL6A1
CAMTA2
BRD3
ARHGEF12
ARHGAP20
Schwalbe
TM
CM
Pericyte
cEndo
VascularEndo
Melanocyte
Schwann
AFF3
ABRA
TMEM136
SPRED2
ME3
HGF
DLC1
DAAM2
CSPG4
BCAS3
ABO
TSPAN18
TRIOBP
TNXB
TMCO1
STOX2
STAG1
SPTBN1
NUP160
NEDD4
MIPOL1
MAPK8IP3
LOX
IL34
HS3ST1
HLA−DQA1
FNDC3B
EXOSC10
EMCN
DLL1
COL4A1
CAPZA1
BTBD3
ANGPTL2
ANGPT2
AFAP1
WBP4
VEGFC
UBE2L3
TRAM2
TRAF3IP2
THSD7A
TCF7L2
TCF4
ST7L
SIX3
SEMA3F
SEC14L1
RPLP2
RPA3
RALB
PRSS23
OXR1
NPAS2
MYOF
MEIS2
MECOM
HACE1
FOXP1
FEZ2
EXOC6
EXOC2
ETS2
CWC25
CRK
CLIC5
CCDC68
BBS9
ADAMTS6
ZNF516
TYR
TSPAN10
TRAPPC3
TEX41
TANGO2
PTPRJ
PTPN1
PKN2
FMNL2
FER
FAM213A
ETS1
CAV2
CAV1
BRE
SEPT9
ZNF652
TRDN
ST3GAL1
SH3PXD2A
GAS7
FOXO1
CADM2
ABCA6
Schwalbe
TM
CM
Pericyte
cEndo
VascularEndo
Melanocyte
Schwann
Expression strength
2
1
0
−1
−2

### Figure S3

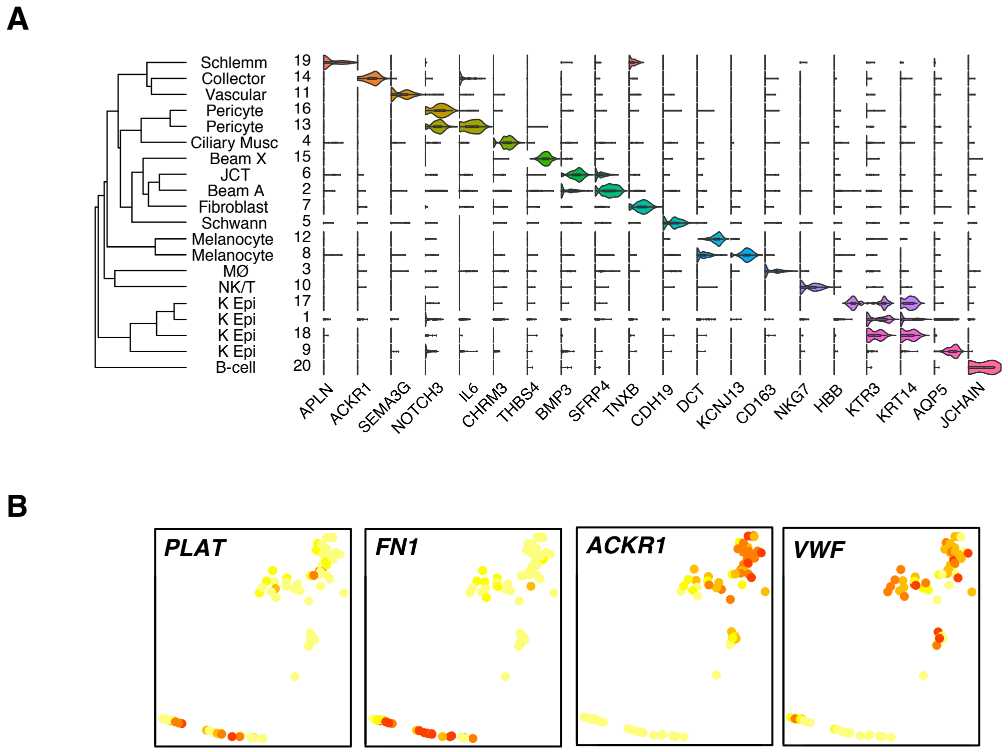
A
Schlemm
Collector
Vascular
Pericyte
Pericyte
Ciliary Musc
Beam X
JCT
Beam A
Fibroblast
Schwann
Melanocyte
Melanocyte
MØ
NK/T
K Epi
K Epi
K Epi
K Epi
B-cell
19
14
11
16
13
4
15
6
2
7
5
12
8
3
10
17
1
18
9
20
APLN
ACKR1
SEMA3G
NOTCH3
IL6
CHRM3
THBS4
BMP3
SFRP4
TNXB
CDH19
DCT
KCNJ13
CD163
NKG7
HBB
KTR3
KRT14
AQP5
JCHAIN
B
PLAT
FN1
ACKR1
VWF

#### Figure S4. Expression Patterns of Selected Genes in Mouse

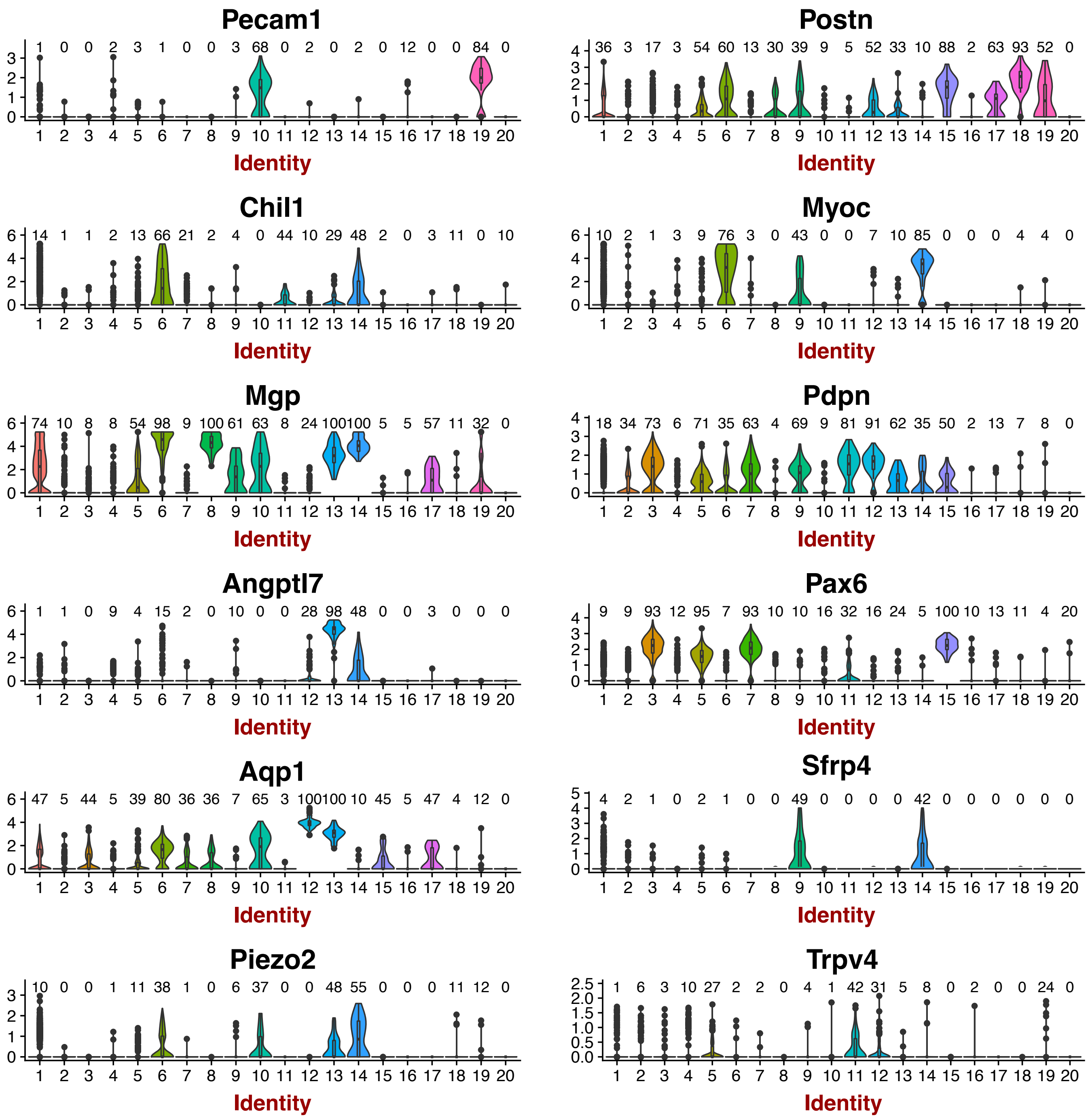

**Table S1. Donor Information**

| Label | Source | 10X | Age | Sex | COD | DTP (Death to processing) | Batch# |
|---|---|---|---|---|---|---|---|
| Pt1 | MGH | V2 | 74 | M | Lung Cancer | 6 | H1TM1 |
| Pt2-RT | MGH | V2 | 78 | M | Metastatic Melanoma to brain | 14 | H2TM1 |
| Pt2-LT | MGH | V2 | 78 | M | Metastatic Melanoma to brain | 14 | H2TM2 |
| Pt3-RT | MGH | V2 | 60 | M | Left tonsillar squamous cell carcinoma metastatic to brain and left orbit | 6.5 | H3TM1 |
| Pt4-LT | MGH | V2 | 64 | M | Diffuse B cell lymphoma spread to thorax and epigastrum | 5 | H4TM1 |
| PT9-LT | MGH | V2 | 53 | F | Interstitial Lung Disease | 5 | H9RimS1 |
| Pt11-RT | MGH | V3 | 65 | M | Metastatic Melanoma | 3 | H11TM1 |

**Table S2. Histological Donor Information**

| Label | Source | Age | Sex | COD | DTP (hr) | In situ | IHC |
|---|---|---|---|---|---|---|---|
| Pt5-LT | MGH | 44 | F | Diffuse B cell lymphoma | 5 | N | LYVE1, CD163, CHRM3, DES |
| Pt6-LT | MGH | 69 | M | Metastatic melanoma | 3 | PPP1R1B, TMEFF2, TFF3, POSTN | CALB2, IL1RL1, CD27 |
| Pt7-LT | MGH | 52 | F | Traumatic brain hemorrhage | 4 | ANGPTL7, CHI3L1 | N |
| Pt10-RT | MGH | 76 | M | Metastatic melanoma | 7 | N | Remnant PDPN and PECAM1 |
| Pt11-RT | MGH | 65 | M | Metastatic melanoma | 3 | RARRES1, CYP1B1 | N |
| HCS6-RT | Lions | 26 | F | Acute respiratory failure | 10 | N | CDH19, DES |
| HCS13-LT | Lions | 22 | F | Multiorgan failure | 10 | N | ACKR1, PECAM1, DES, MLANA, HUD |
| HCS15-RT | Lions | 87 | F | Hemorrhagic shock | 9 | N | PECAM1, ALPL |
| HCS17-RT | Lions | 42 | M | UGIB, cirrhosis | 23.5 | N | PDPN, AQP1 |
| HCS18-LT | Lions | 62 | F | Sepsis | 18 | N | NDUFA4L2, PECAM1 |
| HCS19-RT | Lions | 62 | F | Sepsis | 18 | N | RARRES1, PDPN, FABP4, ADH1B |
